# Molecular dynamics analysis of fast-spreading severe acute respiratory syndrome coronavirus 2 variants and their effects in the interaction with human angiotensin-converting enzyme 2

**DOI:** 10.1101/2021.06.14.448436

**Authors:** Anacleto Silva de Souza, Vitor Martins de Freitas Amorim, Gabriela D A Guardia, Felipe R C dos Santos, Filipe F dos Santos, Robson Francisco de Souza, Guilherme de Araujo Juvenal, Yihua Huang, Pingju Ge, Yinan Jiang, Prajwal Paudel, Henning Ulrich, Pedro A F Galante, Cristiane Rodrigues Guzzo

## Abstract

Severe acute respiratory syndrome coronavirus 2 (SARS-CoV-2) is evolving with mutations in the Spike protein, especially in the receptor-binding domain (RBD). The failure of public health measures to contain the spread of the disease in many countries has given rise to novel viral variants with increased transmissibility. However, key questions about how quickly the variants can spread and whether they can cause a more severe disease remain unclear. Herein, we performed a structural investigation using molecular dynamics simulations and determined dissociation constant (*K*_D_) values using surface plasmon resonance (SPR) assays of three fast-spreading SARS-CoV-2 variants, Alpha, Beta and Gamma ones, as well as genetic factors in the host cells that may be related to the viral infection. Our results suggest that the SARS-CoV-2 variants facilitate their entry into the host cell by moderately increased binding affinities to the human ACE2 receptor, different torsions in hACE2 mediated by RBD variants, and an increased Spike exposure time to proteolytic enzymes. We also found that other host cell aspects, such as gene and isoform expression of key genes for the infection (*ACE2, FURIN* and *TMPRSS2*), may have few contributions to the SARS-CoV-2 variants infectivity. In summary, we concluded that a combination of viral and host cell factors allows SARS-CoV-2 variants to increase their abilities to spread faster than wild-type.

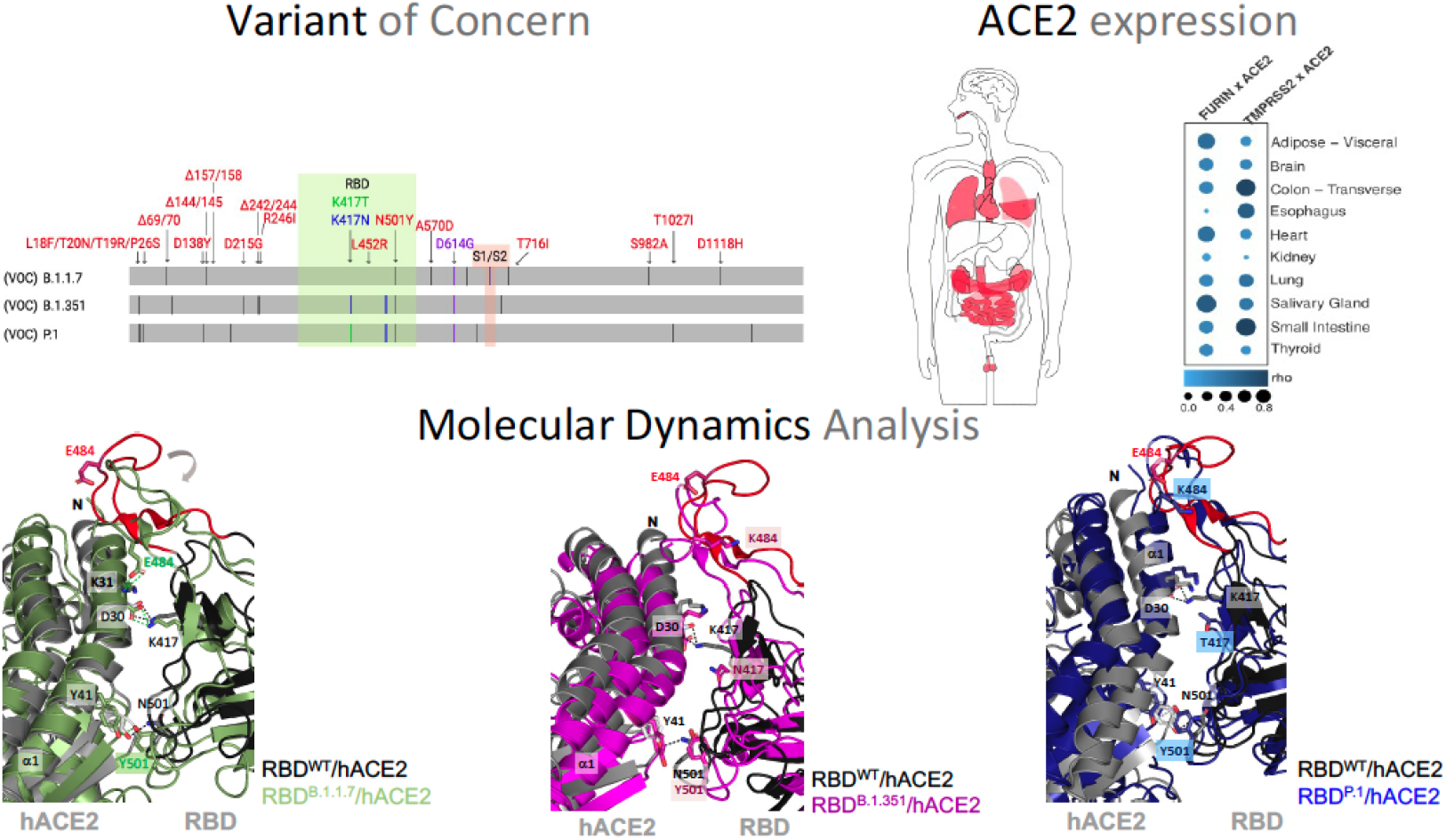

---

We are facing an unprecedented pandemic situation caused by the severe acute respiratory syndrome coronavirus 2 (SARS-CoV-2) ^1,2^, responsible for the death of more than 3,718,683 people and infected more than 170 millions worldwide, as on June 06^th^, 2021^3^. The first outbreak of SARS-CoV-2 was reported in December 2019 at Wuhan, China, causing a severe respiratory syndrome known as Coronavirus Disease 2019 (COVID-19)^3^. The virus surface glycoprotein Spike (S) is composed of two subunits, S1 and S2. S1 has the receptor-binding domain (RBD) that interacts with the human angiotensin-converting enzyme 2 (hACE2) ^4^, while S2 mediates viral and host cellular membrane fusions ^5,6^. The process of virus invasion only occurs after cleavage of two proteolytic sites in the S protein, S1/S2, processed by a furin-like protein convertase (FURIN), and S2’, processed by a transmembrane protease surface serine 2 (TMPRSS2) ^7-10^. A third protease also enhances virus entry, the lysosomal cysteine protease cathepsin L (CTSL) ^10^. These proteolytic processes cleave the S1 subunit and induce drastic conformational changes in the S2 subunit allowing the membrane fusion and release of virus genetic information inside the host cell.

Actually, the world has already gone through two waves, and a third one is imminent with the emergence of novel SARS-CoV-2 variants. According to the World Health Organization (WHO) criteria, a variant of concern (VOC) is defined mainly due to its increased transmissibility, higher virulence or altered clinical characteristics of the disease ^11^. The first VOC reported was in the United Kingdom, also called B.1.1.7 (Alpha variant; Sept 2020), which quickly became prevalent in the British population ^12^. Now, there are three more worldwide spreaded variants: i) B. 1.351, originated in South Africa (Beta variant; May 2020); ii) P.1, originated in Brazil (Gamma variant; Nov 2020); iii) and B. 1.617.2, originated in India (Delta variant; October 2020)^13^. It is evident that these variants have already affected the trajectory of the pandemic in countries where they originated. The SARS-CoV-2 population seems to be still adapting to its human host, since an accumulation of multiple mutations in the S protein are being observed in new variants (**Figure 1a**). Some virus variants have been reported to increase transmissibility, lethality, and to be associated with decreased effectiveness of neutralizing antibodies (**Tables S1** and **S2**) ^14,15^. However, key questions remain about how quickly the variants can spread, their potential to evade immunity and whether they can cause a more severe disease. In order to shed light on these key questions, we performed computational and experimental assays to investigate the changes in binding affinity of three fast-spreading SARS-CoV-2 variants, the Alpha, Beta and Gamma ones, and their cellular entrance, the hACE2. We thus discuss the possible implications of our molecular interaction results to the higher spread and infectivity of the variants.

**Figure 1.**
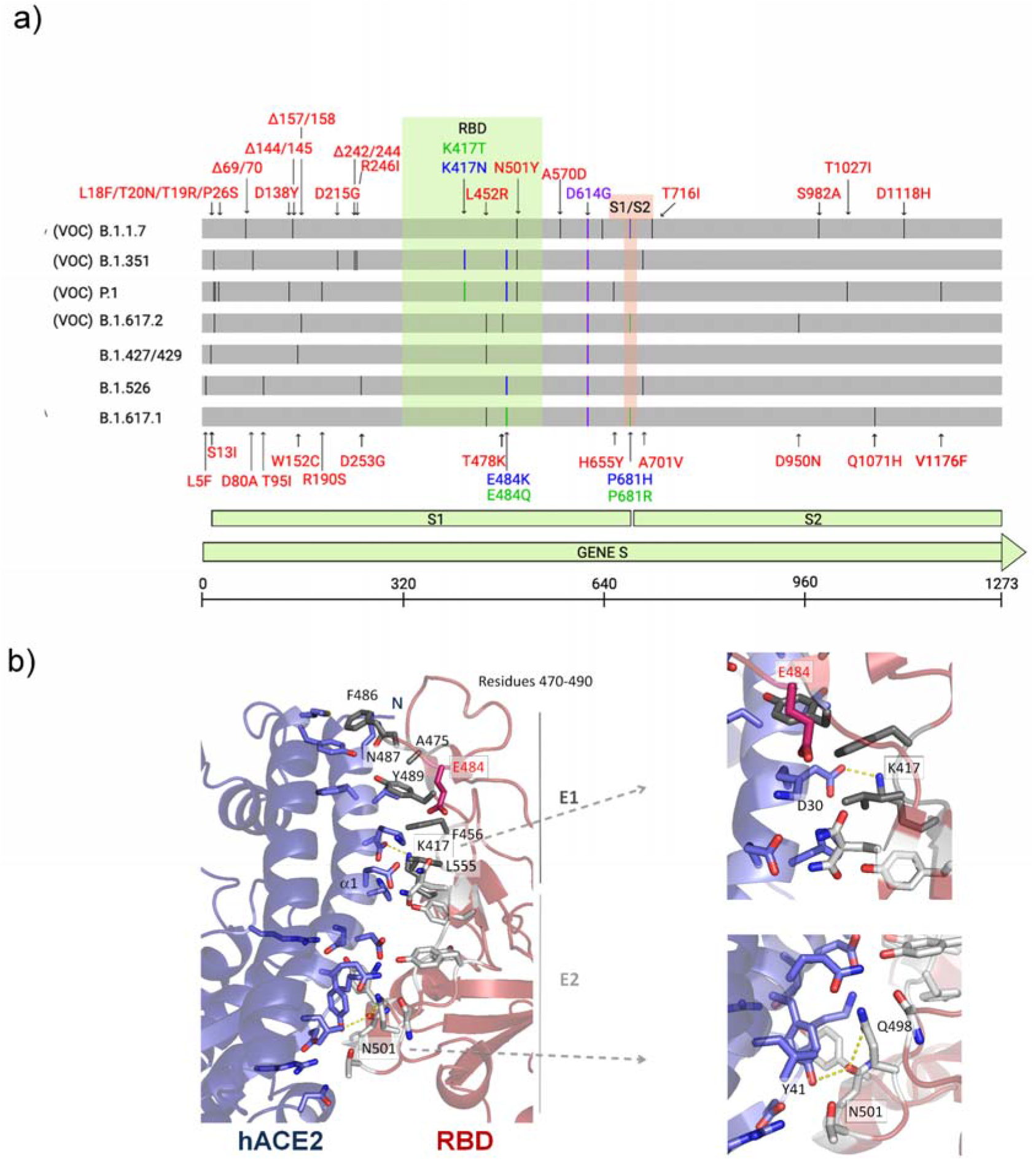
Illustration of mutations in the S protein gene in different variants of SARS-CoV-2. **a)** The RBD (residues 333-526) and the furin cleavage site (S1/S2) (residues 682-685) are colored in light green and blue boxes, respectively. All mutations are represented by black lines and type in red. Mutations that are found to be mutated for different residues are colored in blue and green. The green line shows different mutations in the same location. All SARS-CoV-2 variants shown in the figure have D614G, an advantage mutation for SARS-CoV-2. D614G has enhanced Spike stability and transmission but does not significantly increase binding affinity for hACE2 at 37 0C ^37-39^. VOC: Variant of Concern. **b**) Cartoon representation of the RBD/hACE2 interface complex (PDB ID: 6M0J^4^). RDB presents two different regions, E1 (residues 417, 455–456, and 470–490) and E2 (444–454 and 493-505). The residues mutated in SARS-CoV-2 variants are K417, E487 (does not make part of the interface), and N501. The RBD^B.1.1.7^ (Alpha variant) has the mutation N501Y, the RBD^B.1.351^ (beta variant) has K417N, E484K and N501Y mutations, and the RBD^P.1^ (Gamma variant) has K417T, E484K and N501Y mutations.

In order to evaluate the potential impact of fast-spreading variants B.1.1.7 (RBD^B.1.1.7^), B.1.351 (RBD^B.1.351^), and P.1 (RBD^P.1^) in the interaction with hACE2, we performed molecular dynamics (MD) simulations for 100 ns using RBD (residues 333-526) and hACE2 (residues 19-615) (**Figure 2**). SARS-CoV-2 RDB binds to hACE2 through two different regions, the E1 (residues 417, 455-456, and 470-490) and E2 (444-454 and 493-505) (**Figure 1b**)^16^. While E1 mediates more hydrophobic interactions, E2 mediates mostly polar interactions with hACE2. RBD^P.1^ has mutations in K417T, E484K and N501Y; RBD^B.1.351^ has mutations in K417N, E484K and N501Y; and RBD^B.1.1.7^ has only the N501Y mutation. K417 and N501 are in the protein complex interface, whereas E484 is located in a mobile loop distant from the interface (**Figure 1b**). From the reference^17^, we verified that mutations in the protein interface cause a hydrophobicity gain (N501Y of +2.2 kcal/mol, K417T of +4.6 kcal/mol, K417N of +0.4 kcal/mol) that may result in an enhancement in complex stability.

**Figure 2.**
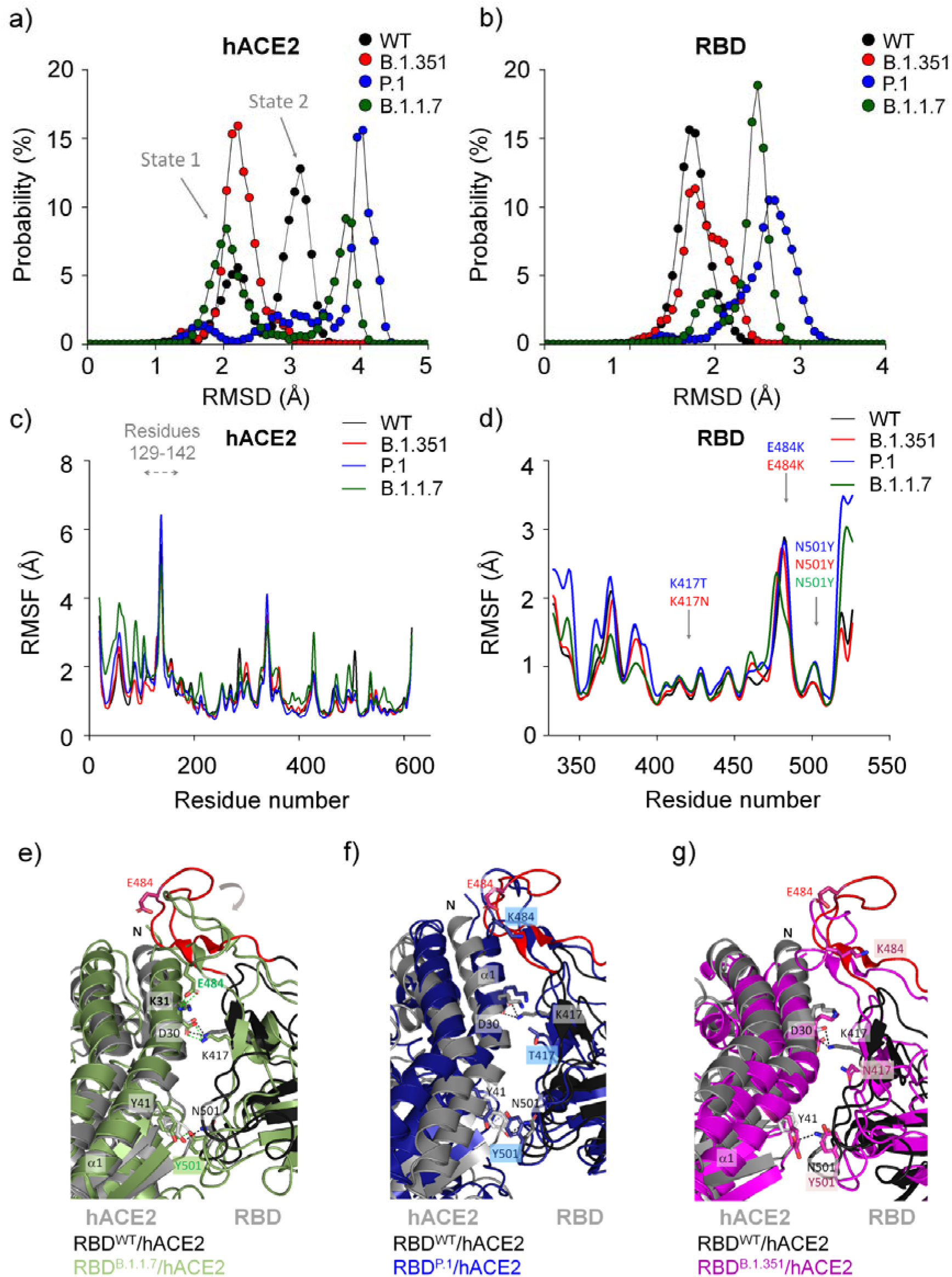
Structural aspects obtained by molecular dynamics simulations of different SARS-CoV-2 RBD variants while interacting with hACE2. Backbone root-mean-square deviation (RMSD) probability distributions are shown for **a)**hACE2 and **b**) RBD. Backbone root-mean-square fluctuation (RMSF) are shown for **c**) hACE2 and **d**) RBD. The panels **e-g** show different conformational changes of each RBD^variant^/hACE2 complex in relation to RBD^WT^/hACE2 complex. Note that main structural differences are localized in the loop 474-488 caused by N501Y mutation in RBD^B.1.1.7^ (panel e) that is partially recovered in RBD^P.1^ (panel f) and RBD^B.1.351^ (panel g).

Molecular dynamics (MD) simulations analysis showed that hACE2 presented different backbone root-mean-square deviation (RMSD) and backbone root-meansquare fluctuation (RMSF) profiles in the presence of different RBD variants, RBD^B.1.1.7^, RBD^B.1.351^ and, RBD^P.1^ (RBD^variants^) (**Figures 2** and **S2**). Consistent with our previous study ^16^, hACE2 sampled in two major conformational states (state 1 of RMSD of ~2.4 Å and state 2 of ~3.2 Å, **Figures 2a**), characterized by a significant motion of residues 129-142 located distant from the interface (**Figure 2c**). In the interaction with RBD^WT^ and RBD^B.1.1.7^, hACE2 presented RMSD^WT^ values of ~2.2 and ~3.1 Å and RMSD^B.1.1.7^ values of ~2.0 and ~3.8 Å, both sampling two conformational states. In contrast, in the interaction with RBD^P.1^, hACE2 sampled three states, presenting RMSD values of ~1.8, ~2.8 and ~4.0 Å (**Figure 2a**). While interacting with RBD^B.1.351^, hACE2 visits one major state (**Figure 2a**). Probability distributions of the backbone RMSD along the MD trajectory showed that the RBD mutants sampled a broader conformational ensemble in comparison to RBD^WT^ (**Figure 2b**). Interestingly, hACE2 presents significant changes in solvent-accessible surface area (SASA) in complex with different RBD variants, while RBDs share similar values (**Figure S4**). This observation suggests that RBDs induce different conformational states in the hACE2.

Curiously, the RMSF profile is slightly different for the RBD^B.1.1.7^/hACE2 complex in relation to the others: while RBD^B.1.1.7^ decreases its RMSF values for residues 350-400 (distance of the interface), hACE2 increases its RMSF values for residues 19-140, which are located in the interface (**Figures 2d**). The K417 mutation does not significantly affect its RMSF values, yet mutation in N501Y slightly increases its flexibility in the RBD^P.1^ and RBD^B.1.1.7^ and it is restored in the RBD^B.1.351^. Intriguingly, N501Y mutation in RBD^B.1.1.7^ causes a loss of flexibility in the residues 480-486, which is restored by E484K mutation in RBD^P.1^ and RBD^B.1.351^ (**Figure 2d-g**). Based on the analysis of the snapshots of RBD/hACE2 complexes, it is evident that the mutation of N501Y in RBD^B.1.1.7^ causes a significant folding in the loop containing residues 474-488 (more mobile region of the RBD), favoring the formation of a hydrogen bonding between E484^RBD^ and K31^hACE2^ (**Figure 2e** and **Table S3**). In agreement, a similar folding is also observed in the cryo-electron microscopy of the trimeric Spike structure of the B.1.1.7 variant (**Figure S3**)^18^. Our analyses of hydrogen bonding occupancies revealed absence of hydrogen bonding interaction between E484^RBD^ and K31^hACE2^ in the other complexes (**Figure 2f-g; Table S3**). Additionally, RBD^P.1^ and RBD^B.1.351^ have an E484K mutation, causing a charge-repelling impact with K31^hACE2^. These variants also present the mutations K417T and K417N, respectively. The K417 and E484 mutations seem to reverse the effect of the N501Y mutation in the loop of residues 474-488, restoring a more original conformational state (**Figures 2c-g** and **S2a-b**).

We also calculated the covariance matrix of the spatial displacements of C_α_ atoms to study internal motions of the hACE2/RBD^WT^ complex and compared them with different SARS-CoV-2 RBD mutants (**Figures 3** and **S5**; **Tables S4** and **S5**). The sets of intermolecular and intramolecular covariant pairs changed between different SARS-CoV-2 RBD mutants and RBD^WT^ (**Table S4** and **S5**). Both correlated and anticorrelated pairs were affected in E1 and E2 regions of RBD variants, mostly in RBD^B.1.1.7^, whereas the other two mutants tend to show similar patterns to RBD^WT^ (**Figure 3**). We also calculated the principal components using MD trajectories to provide the main structural insights of the RBD/hACE2 complexes from eigenvalues and orthogonal eigenvectors (**Figures 4** and **S6**). These orthogonal eigenvectors are associated with the maximum covariance between residue pairs. Thus, we observed 100 snapshots corresponding to MD trajectories obtained by the first eigenvalue of the RBD^WT^/hACE2 and RBD^variants^/hACE2 complexes (**Figure 4**). The movement of the RBD and hACE are different in each analyzed complex, demonstrating that RBD induces different torsions in the hACE2 (**Figure 4** and **supplementary movies S1** and **S2**). In this regard, we hypothesized that observed torsions in hACE2 may be important for mediating efficiently structural transition of the Spike protein from the pre-fusion to the fusion intermediate state ^19^, facilitating the virus entry into the host cell. Therefore, the different conformational changes of RBD^variants^ interacting with hACE2 could facilitate the transition process from prefusion to postfusion state.

**Figure 3.**
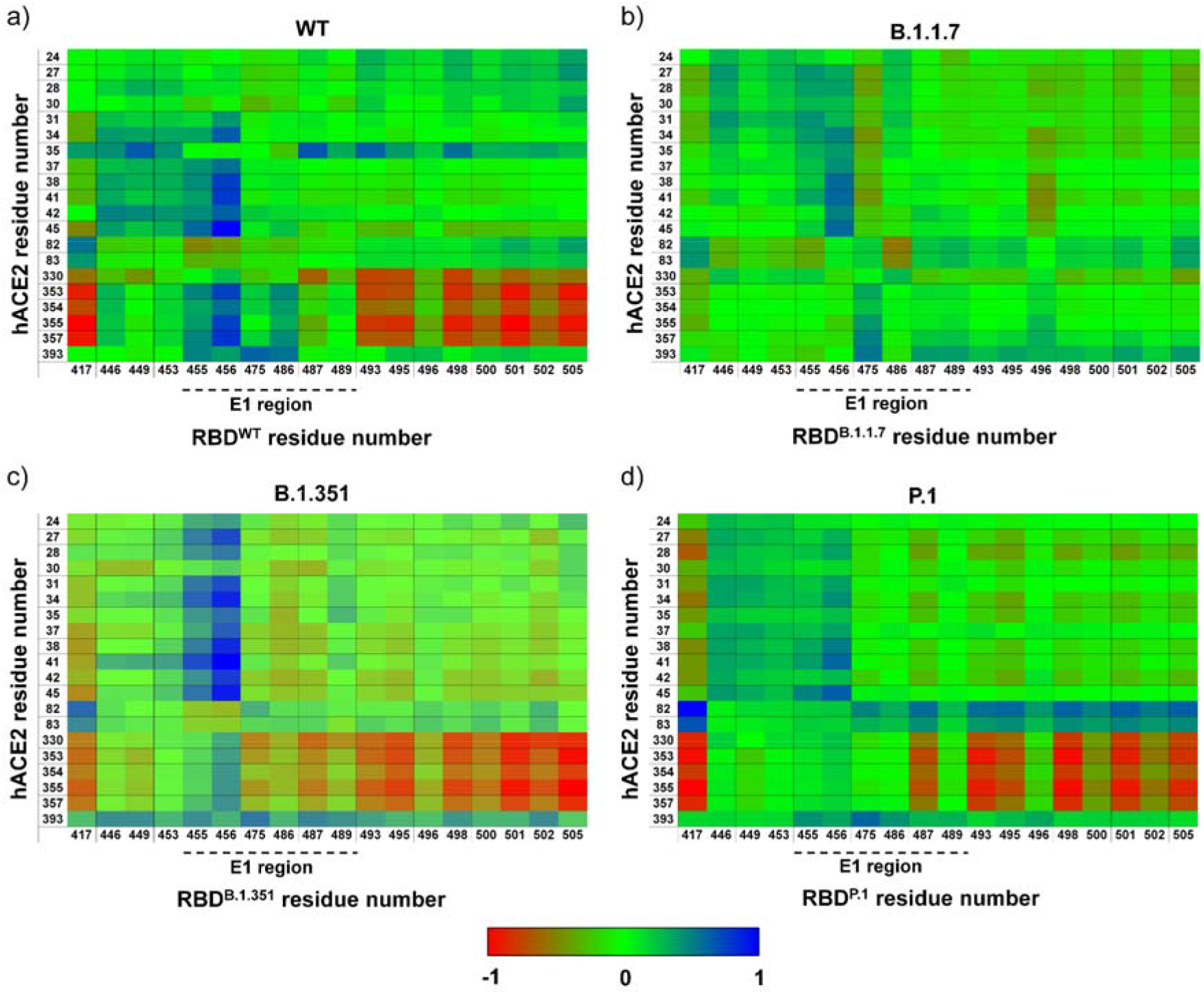
Covariant matrix obtained by MD trajectories of RBD and its variants interacting with hACE2. Painels **a-d)** show heat maps representing correlation matrix of the interface residues. Anti-correlated and correlated pairs are colored in blue and red colors, respectively. Note that correlated and anticorrelated pairs are affected in E1 and E2 regions of RBD variants, mostly in RBD^B.1.1.7^. Heat maps of the RBD^B.1.351^ and RBD^P.1^ have similar patterns to RBD^WT^.

**Figure 4.**
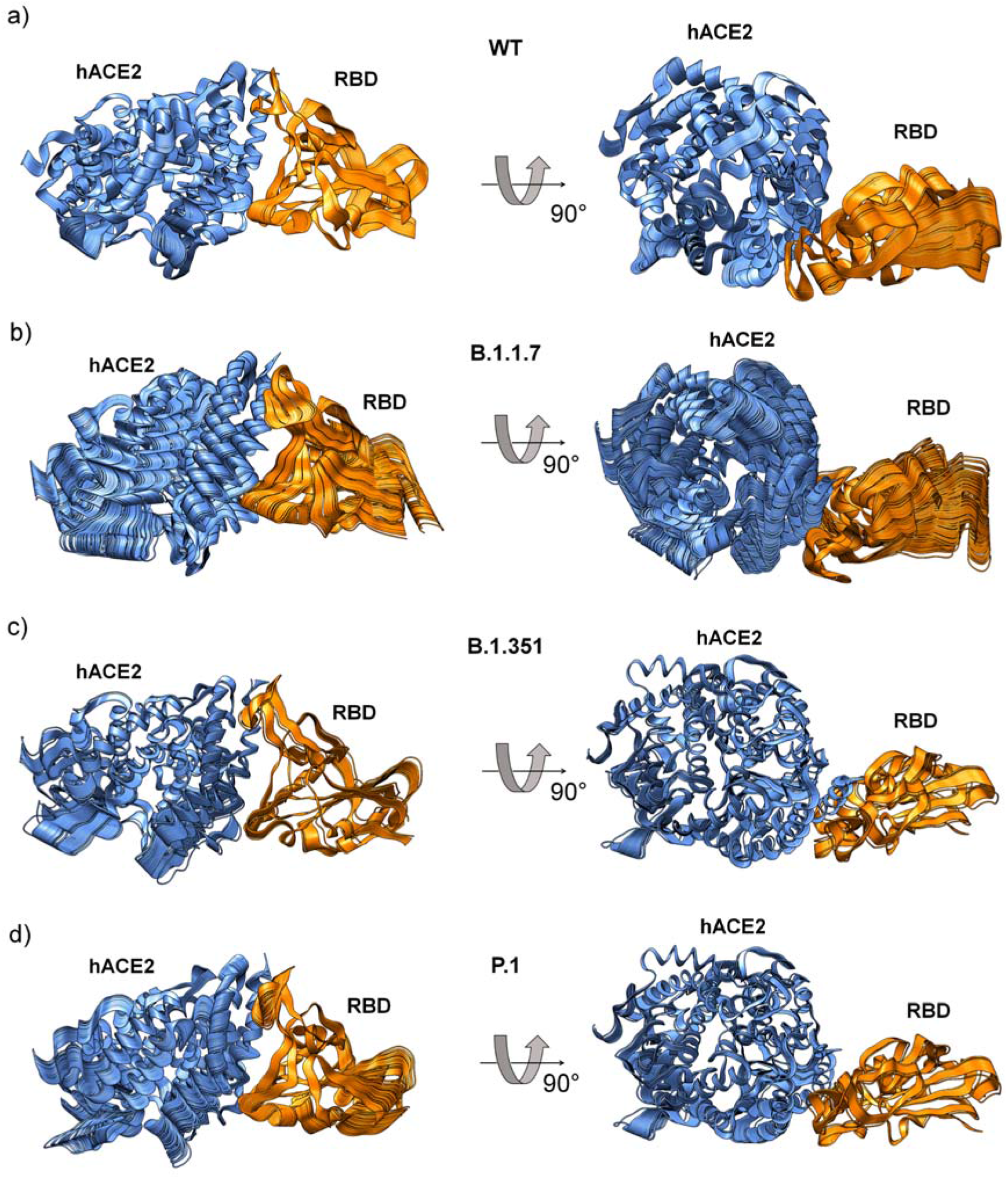
Molecular dynamics trajectories obtained by principal component analysis. Principal component analysis using MD trajectories of hACE2 in complex with **a**) RBD^WT^, **b**) RBD^B.1.1.7^, **c**) RBD^B.1.351^ and **d**) RBD^P.1^. Each painel has 100 snapshots corresponding to MD trajectories obtained from the first component. The movements of the RBD variants induce different torsions in the hACE2 (**supplementary movies S1** and **S2**).

We then investigated the contributions of the RBD^variants^ in the hACE2 complex formation by calculating the binding free energy using Molecular Mechanics combined with Poisson-Boltzmann Surface Area (MM-PBSA). The binding free energy ratio between RBD^WT^/hACE2 and RBD^variants^/hACE2 complexes presented values ranging from ~1.0-fold to ~1.4-fold (**Table S6** and **Figure S6**). Most contributions for slight increasing binding affinity for RBD^variants^/hACE2 complexes are localized on the interaction interface residues (**Figure S7**). We also performed surface plasmon resonance (SPR) assays to measure the dissociation constant (*K*_D_)values using hACE2 and different constructions of trimeric Spike and RBD variants (**Tables 1** and **S7; Figures S8** and **S9**). RBD^variants^ bind hACE2 with *K*_D_ values ranging from 1.2 to 3.3 nM, showing that they have more binding affinity for hACE2 than RBD^WT^ (*K*_D_ = 10.3 nM). Trimeric Spike variants also increased their binding affinities for hACE2, ranging from 1.1 to 1.6 nM, when compared to trimeric Spike^WT^ (*K*_D_ = 6.4 nM). Our data are consistent with other studies showing that variants of the SARS-CoV-2 Spike protein have greater affinity for hACE2 than wild-type (**Table S8**) ^20-24^. Although there are only slight differences in the *K*_D_ values, significant differences in *k*_a_ and *k*_d_ rates point to altered binding kinetics (**Table 1** and **S7**). Decreases in the *k*_d_ rates in these variants indicate a longer permanence of the Spike/hACE2 complex when compared to the wild-type, which is necessary for successful cleavage of the Spike protein and cell invasion by the virus. Decreased *k*_d_ rates agree with an enhanced goodness of fit for virus-ACE2 protein interactions.

**Table 1.** Kinetic parameters obtained from surface plasmon resonance (SPR) assays. Equilibrium dissociation constants (*K*_D_) calculated for RBD and its variants in the complex with dimeric hACE2 protein. Experimental *K*_D_ values were also measured using a trimeric Spike protein and its variants for interacting with hACE2. The mutations in each Spike constructions are shown in parentheses: RBD^B.1.1.7^ (N501Y); RBD^B.1.351^ (K417N, E484K N501Y); RBD^P.1^ (K417T, E484K N501Y); Spike trimer^B.1.1.7^ (H69-V70del, Y144del, N501Y, A570D, D614G, P681H, T716I, S982A, D1118H); Spike trimer^B.1.351^ (L18F, D80A, D215G, L242-A243-L244del, R246I, K417N, E484K, N501Y, D614G, A701V); Spike trimer^P.1^ (L18F, T20N, P26S, D138Y, R190S, K417T, E484K, N501Y, D614G, H655Y, T1027I, V1176F). In Spike trimer constructions the proline substitutions (F817P, A892P, A899P, A942P, K986P, V987P) were introduced to stabilize the trimeric prefusion state of SARS-CoV-2 Spike protein and alanine substitutions (R683A and R685A) were introduced to abolish the furin cleavage site.

| Variant | $k_a$ ($10^6 \text{ M}^{-1} \text{ s}^{-1}$) | $k_d$ ($10^{-4} \text{ s}^{-1}$) | $K_D$ (nM)* |
| --- | --- | --- | --- |
| RBD<sup>WT</sup> | 0.9 | 91.6 | 10.3 |
| RBD<sup>B.1.1.7</sup> | 1.3 | 15.5 | 1.2 |
| RBD<sup>B.1.351</sup> | 1.2 | 39.4 | 3.3 |
| RBD<sup>P.1</sup> | 1.3 | 28.8 | 2.2 |
| Spike trimer<sup>WT</sup> | 0.09 | 5.8 | 6.4 |
| Spike trimer<sup>B.1.1.7</sup> | 0.1 | 1.7 | 1.6 |
| Spike trimer<sup>B.1.351</sup> | 0.3 | 3.0 | 1.1 |
| Spike trimer<sup>P.1</sup> | 0.2 | 3.0 | 1.8 |

\* $K_D = k_d/k_a$

Taken together, even a small gain in binding affinity of the mutants may enhance infection, facilitating hACE2/Spike complex formation, necessary for membrane fusion and virus entry^25^. It is also possible that actions of allosteric domains in the Spike protein, priming Spike/hACE2 binding ^26,27^, benefit from the mutations in the SARS-CoV-2 variants. It is relevant to mention that Spike/hACE2 interactions and the binding affinity for complex formation also depends on the microenvironment. For instance, the presence of heparin or heparan sulfate ^28,29^, among others, so far not well-studied, positively or negatively acting allosteric factors that might provide an explanation for the differences between *K*_D_ values from our experimental kinetic data and other studies. In addition to any favoring of Spike/hACE2 complex formation, mutants may facilitate FURIN binding and promote cleavage, as shown for the SARS-CoV-2 Spike mutant protein D614G ^30^. Further biological factors are probably associated with the increased infectivity of SARS-CoV-2 variants. It has been reported that D614G mutation can increase the density of Spike on the surface of the virion due to a decrease in the premature loss of the S1 subunit ^31^. It is known that P.1, B.1.1.7 and B.1.351 variants have lower affinity for neutralizing antibodies, which may be a way for the virus to escape the immune system of an infected individual ^21,32–34^. Additionally, factors related to the expression of key genes and their variants for SARS-CoV-2 infections may be critical.

We investigated whether expression profiles of three essential genes for the viral entrance into the human cells, *ACE2, TMPRSS2, and FURIN* may be determinative for infection rates. We analyzed approximately 23,000 samples (~1,000 individuals) from 32 healthy tissues and, to gain more insight about their expression patterns, we also sectioned them in gender, donors’ age (young < 40; old > 60 years old) and *ACE2* splicing isoforms (**Figure 5**). *ACE2* has a high or middle expression in organs of respiratory tract (lung), cardiovascular (heart), digestive (colon and small intestine), endocrine (testis, thyroid), adipose tissue, breast and kidney (**Figure 5a**). *TMPRSS2* has a similar expression pattern. Notably, both genes are low expressed in the nervous system (**Figure 5a** and **Figure S10**). Conversely, *FURIN* is highly expressed in almost all human tissues (**Figure S11**). **Figure 5b** summarizes organs presenting high expression of *ACE2* and *TMPRSS2* or *FURIN* (red marked). Most of them are already known to be critical organs for the disease (*e.g*., lung and heart). Interestingly, a new ACE2 splicing isoform was recently reported ^35,36^, ^35,36^ which lacks the S1 RBD interaction region, is induced by interferons but has no metallopeptidase activity. Then, we investigated the expression of this isoform (*dACE2*), as well as the full-lenght (*fACE2*) and other *ACE2* alternative isoforms (*pACE2*) (**Figure S12**). **Figure 5c** shows that the fulllength isoform, which contains receptor regions that interact with S RBD, is majorly expressed in all human tissues. Next, we also investigated the expression of these three genes by gender and age (young < 40; old > 60 years old). We did not find significant differential expression by age or gender for *ACE2* (**Figure 5d**), *TMPRSS2* (**Figure S13**) or FURIN (**Figure S14**). Finally, we found a high expression correlation between *ACE2* and the proteases, *FURIN* and *TMPRSS2*, suggesting a functional synergy in organs usually affected by the SARS-CoV-2 (**Figure 5e**). In summary, we confirmed that *ACE2* and either proteases, *FURIN* or *TMPRSS2*, are co-expressed in organs already described as affected by SARS-CoV-2, such as the respiratory, cardiovascular, digestive, and some endocrine organs. Using these expression data (**Materials and Methods**, in **supporting information**), no significant expression differences were found for these genes in terms of gender, age and *ACE2* splicing isoforms.

**Figure 5.**
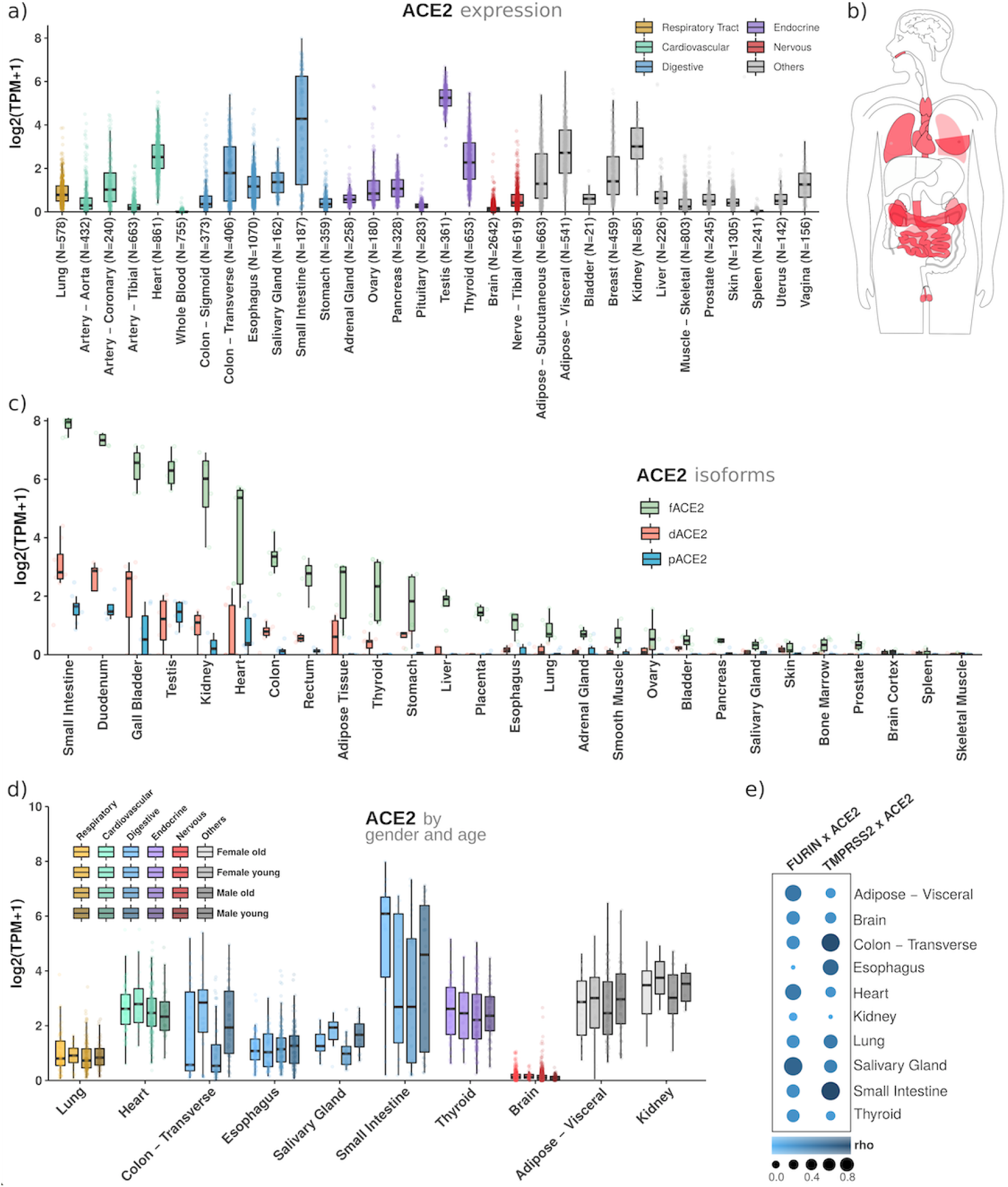
*ACE2* is expressed in critical tissues for the disease, its splicing isoforms are less expressed than its full-length sequence, and there is no significant expression difference between gender or age for this gene. **a)** *ACE2* gene expression profile in 32 human healthy tissues. **b)** Schematic representation of organs showing higher expression of *ACE2* and *TMPRSS2* or *FURIN* genes. **c)** Expression profile of *ACE2* splicing isoforms. **d)** *ACE2* expression profile segmented by age (young < 40; old > 60 years old) and gender. **e)** Expression correlation (Rho = spearman coefficient) between *ACE2* and *FURIN* or *ACE2* and *TMPRSS2*.

Overall, our computational and experimental data show that the SARS-CoV-2 variants moderately increase their binding affinities to hACE2, which is not enough to explain the increase of transmission and lethality caused by the virus. We suggested that decrease of *k*_d_ rates values could positively influence the trimeric Spike cleavage, by increasing the exposure time of its cleavage site to proteolytic enzymes. Furthermore, SARS-CoV-2 variants modulate different torsions in hACE2, which may efficiently mediate structural transition of the Spike protein from the prefusion to the fusion intermediate state^19^, facilitating the virus entry into the host cell. The individual genetic background, such as *ACE2, FURIN* and *TMPRSS2* polymorphisms and expression patterns, among other factors, will together with circulating and upcoming SARS-CoV-2 variants determine the course and duration of the pandemics. Therefore, combining all these components allows SARS-CoV-2 variants to increase their abilities to spread faster than wild-type.

## Acknowledgment

The authors acknowledge the National Council for Scientific and Technological Development (CNPq), the Coordination for the Improvement of Higher Education Personnel (CAPES, grant 88887.374931/2019-00 and 88887.620198/2021-00, Coordenação de Aperfeiçoamento de Pessoal de Nível Superior - Finance Code 01), and the São Paulo Research Foundation (FAPESP, grants 2019/00195-2, 2020/04680-0, 2016/09047-8, 2018/07366-4, 2018/15579-8, 2017/19541-2, 2017/18246-7, 2017/17636-6, 2020/14158-9 and 2020/06091-1), Brazil, for financial support.

