## supplemental information for "Molecular dynamics analysis of fast-spreading severe acute respiratory syndrome coronavirus 2 variants and their effects in the interaction with human angiotensin-converting enzyme 2"

**Running title: Molecular dynamics analysis of fast-spreading SARS-CoV-2 variants and their effects in the interaction with hACE2**

To whom correspondence should be addressed:

Cristiane R. Guzzo, Ph.D, Department of Microbiology, Institute of Biomedical Sciences, University of São Paulo, Av. Prof. Lineu Prestes, 1374, Cidade Universitária, 5508-900, São Paulo/SP, Brazil, +55 11 3091-7298;

**Keywords:** SARS-CoV-2 variants, binding affinity, molecular dynamics, transcriptome analysis, SPR assay, COVID-19.

**Materials and Methods** **1**
Molecular dynamics (MD) simulations 2
Molecular mechanics combined with Poisson-Boltzmann surface area (MM-PBSA) 3
Protein-protein binding assays using surface plasmon resonance (SPR) 3
Genes and splicing isoforms expression analyses 5

**Supplementary text** **6**
Solvent-accessible surface area (SASA) analysis 6
Molecular mechanics combined with Poisson-Boltzmann surface area (MM-PBSA) studies 6
Hydrogen bonding occupancy analysis 7
Cross-correlation analysis 8

**Figures S1-S14.** **9**

**Tables S1-S7.** **23**

### Materials and Methods

#### Molecular dynamics (MD) simulations

The initial coordinates for RBD/hhACE2 complex were obtained from the Protein Data Bank (PDB ID 6M0J, resolution 2.45 Å)[1] and prepared using the UCSF Chimera[2] by removing co-crystallized hetero groups (except $Zn^{2+}$) and water molecules. RBD mutations were obtained from a site-directed mutation in wild-type RBD using the Maestro software (academic v. 2020-1) [3]. From the Ade1 structure, protonation states of ionizable residues were computed in an aqueous implicitly environment at pH 7.0 from the Maestro software academic v. 2020-1 [3] using PROPKA module [4]. In all protein complexes, then, all glutamic and aspartic residues were represented as unprotonated; all glutamic residues were kept with neutral charge; arginine and lysine residues were assumed with a positive charge; the N- and C-terminal were converted to charged groups. In hACE2, H34, H195, H345, H374, H378, and H417 were designed as a δ-tautomer; H228, H239, H241, H265, H373, H401, H505, H535 and H540 were modeled as an ε-tautomer. In RBD, H519 was designed as a δ-tautomer. The system conditions were prepared using GROMACS v. 5.1.4[5–8] with the OPLS-AA force field[9]. We used the Swiss-Param web-based service to build the $Zn^{2+}$ topology[10]. All systems were explicitly solvated with *TIP3P* water models in a triclinic box (81.06 X 91.06 X 135.02 Å³), neutralized keeping NaCl concentration of 150 mM and minimized until a maximum force of 10.0 $kJ.mol^{-1}$ or 5,000 steps. The systems were consecutively equilibrated in isothermal-isochoric (NVT), To equilibrate the system, it was relaxed during a 5000 ps annealing, in which the temperature was increased from 300 to 330 K each 500 ns, and isothermal-isobaric (1 bar; NpT) ensembles at 310 K for 1000 ps. MD simulations in the NpT ensemble were carried out for 100 ns in a periodic cubic box considering the minimum distance of 10 Å between any protein atom and box walls. The $Zn^{2+}$ positions were restricted throughout the simulations. Backbone root-mean-square deviation (RMSD), and backbone root-mean-square fluctuation (RMSF) calculations were performed using GROMACS. Hydrogen bonding occupancies (%) were calculated in virtual molecular dynamics (VMD; v. 1.9.1).

Analysis of $C_\alpha$ cross-correlated displacements were performed using the R-based package Bio3d[11]. The covariances ranged from -1 (anti-correlated motions) to 1 (correlated motions). Principal component analysis and SASA were calculated using the GROMACS package. Using molecular dynamics simulation trajectories, we determined the average structure representing backbone hACE (2373 atoms) and backbone RBD (1791 atoms) and calculated correlation matrix fitting non-mass weighted. From the calculated correlation matrix, we determined eigenvalues diagonalizing the 7119x7119 covariance matrix (representing backbone hACE2) and 5373x5373 covariance matrix (representing backbone RBD). The first and second eigenvalues are shown in **Table S9**. From the PC1, we observed 100 snapshots corresponding to MD trajectories obtained from the first component (**supplementary movies S1** and **S2**). We used first and second components to verify trajectory scores of hACE2 and also $RBD^{WT}$ and their variants (**Figure S6**).

**Molecular mechanics combined with Poisson-Boltzmann surface area (MM-PBSA)**

The molecular mechanics combined with the Poisson-Boltzmann surface area (MM-PBSA) was calculated using the Kumari method[12], which approximates the solvation contribution to a continuous solvent model and approximates ΔG to a thermodynamic cycle by modeling a continuous solvent in the Poisson-Boltzmann area. The charges, radius and concentration of positive ions were +1, 0.95 Å and 0.15 M, respectively. The charges, radius and concentration of negative ions were -1, 1.81 Å and 0.15 M, respectively. The dielectric constant of the solute, solvent and vacuum was 32, 80 and 1, respectively. The radius of the solvent probe was 1.4 Å. The method for mapping the grid was sp14 and the model used to build the dielectric and ionic limits was smol. In the free-energy calculation, we considered for the non-polar solvent, the surface tension, probe radius and energy cut-off of 0.023 $kJ.mol^{-1}.Å^{-2}$, 1.4 Å, 3.85 $kJ.mol^{-1}$, respectively. Non-polar attractive contribution term was determined by the probe radius, the density of solvent in the grid and the cubic grid spacing for the calculation of the volume integrals of 1.20 Å, 0.03 $Å^3$ and 0, 7 Å, respectively. The time interval analyzed was 1 ns.

**Protein-protein binding assays using surface plasmon resonance (SPR)**

The experimental $K_D$ ($M^{-1}$) of the SARS-CoV-2 Spike/hACE2 complex were obtained by surface plasmon resonance (SPR) assays using a Biacore 8K system (GE Healthcare). The assays as well as protein cloning, expression and purification were performed by the ACROBiosystems company. SARS-CoV-2 Spike constructions are fused to a 10 polyhistidine tag at the C-terminus of the recombinant proteins. Four different constructions of the Spike RBD as well as four constructions of recombinant Spike trimers were used in SPR assays: the $RBD^{WT}$ (residues 319 to 537, SPD-C52H3, ACROBiosystems); the $RBD^{B.1.1.7}$ (N501Y, residues 319 to 537, SPD-C52Hn, ACROBiosystems); the $RBD^{B.1.351}$ (K417N, E484K N501Y, residues 319 to 537; SPD-C52Hp, ACROBiosystems); the $RBD^{P.1}$ (K417T, E484K N501Y, residues 319 to 537; SPD-C52Hr, ACROBiosystems); Spike $trimer^{WT}$ (residues 16 to 1213 with the following mutations at: R683A, R685A, F817P, A892P, A899P, A942P, K986P, V987P; SPN-C52H9, ACROBiosystems); Spike $trimer^{B.1.1.7}$ (residues 16 to 1213 with the following mutations at: R683A, R685A, F817P, A892P, A899P, A942P, K986P, V987P and additional mutations at: H69-V70del, Y144del, N501Y, A570D, D614G, P681H, T716I, S982A, D1118H; SPN-C52H6, ACROBiosystems); Spike $trimer^{B.1.351}$ (residues 16 to 1213 with the following mutations at: R683A, R685A, F817P, A892P, A899P, A942P, K986P, V987P and additional mutations at: L18F, D80A, D215G, L242-A243-L244del, R246I, K417N, E484K, N501Y, D614G, A701V; SPN-C52Hk, ACROBiosystems); Spike $trimer^{P.1}$ (residues 16 to 1213 with the following mutations at: R683A, R685A, F817P, A892P, A899P, A942P, K986P, V987P and additional mutations at: L18F, T20N, P26S, D138Y, R190S, K417T, E484K, N501Y, D614G, H655Y, T1027I, V1176F; SPN-C52Hg, ACROBiosystems). In Spike trimer constructions the proline substitutions (F817P, A892P, A899P, A942P, K986P, V987P) were introduced to stabilize the trimeric prefusion state of SARS-CoV-2 Spike protein and alanine substitutions (R683A and R685A) were introduced to abolish the furin cleavage site. The human ACE2 (residues from 18 - 740) is fused to a Fc tag at the C-terminus of the recombinant protein (MW 110 kDa; AC2-H5257, ACROBiosystems), in which hACE2 is in a dimer form.

Anti-Human IgG (Fc) antibody was diluted at 25 μg/mL in an immobilization buffer (10 mM Sodium Acetate pH 5.0 present in GE Human Antibody Capture Kit, GE), to be covalently immobilized to a CM5 sensor chip (GE Healthcare) via their amine groups using the human antibody capture kit (29234600, GE Healthcare). Immobilization processes were performed using a flow rate of Anti-Human IgG (Fc) antibody of 10 μL.min$^{-1}$ during 360 s, where obtained response unit (RU) signals were about 9,000-14,000. Human ACE2 with a C-terminal Fc tag was diluted in a running buffer (10 mM HEPES, pH 7.4, 3 mM EDTA, 0.005% Tween-20, 150 mM NaCl). The RU signal obtained was about 150 in the capture hACE2 protein-ligand. After that, serial dilutions of purified recombinant Spike proteins were injected ranging in concentrations (as shown in **Table S7**). Spike proteins were injected at a flow rate of 30 μL.min$^{-1}$ during association (120 s) and dissociation (300 s). The chip was regenerated using a regeneration buffer (3M magnesium chloride, present in GE Human Antibody Capture Kit, GE) at a flow rate of 20 μL.min$^{-1}$ in 30 s. The resulting data were fitted to a 1:1 binding model using Biacore Evaluation Software (GE Healthcare).

**Genes and splicing isoforms expression analyses**

To investigate the expression profiles of *ACE2*, *FURIN* and *TMPRSS2*, we gathered Transcripts Per Million (TPM)-normalized gene expression data of 23,000 samples from 32 healthy tissues directly at The Genotype-Tissue Expression (GTEx) portal (https://gtexportal.org/). Subject phenotype information (gender and age) of all analyzed individuals were also collected at the GTEx portal and used for posterior data stratification. To investigate isoform-based expression levels of *ACE2*, we downloaded RNA sequencing datasets from 163 samples (27 tissues) at ENA projects (accession IDs: PRJEB4337; PRJEB6971). To obtain TPM-normalized expression values, the datasets were processed using Kallisto [13] with GENCODE (v37) as reference to the human transcriptome. Boxplots and correlation plots were generated using local R scripts. Alignment of translated transcripts was performed through clustal W [14] using *ACE2* transcripts from GENCODE (v37) and ORFfinder [15].

#### Supplementary text

##### Solvent-accessible surface area (SASA) analysis

Such conformational changes affect the solvent-accessible surface area (SASA) of hhACE2 while interacting with $RBD^{WT}$ and its variants (**Figure S4**). In this regard, $RBD^{WT}$ and $RBD^{B.1.351}$ keep SASA values of ~273 and 270 nm², respectively (**Figure S4a,c**). Interacting with $RBD^{B.1.1.7}$, hhACE2 decreases SASA value from ~278 to 263 nm² but when it interacts with $RBD^{P.1}$, SASA value changes from 270 to 260 nm² (**Figure S4a,c**). In general, RBD variants and WT did not present significant differences in SASA along MD trajectory, presenting values ranging between 107.0 and 109.2 $nm^2$ (**Figure S4b,d**). Taken together, SASA values are associated with conformational states of hACE2 due to closing of its active site mediated by RBD and its variants.

##### Molecular mechanics combined with Poisson-Boltzmann surface area (MM-PBSA) studies

We also compared the binding free energies between Spike RBD of the different B.1.1.7, B.1.351 and P.1 variants with $RBD^{WT}$, using molecular mechanics combined with Poisson-Boltzmann surface area (MM-PBSA) (**Figure S7** and **Table S6**). Our results have shown that $RBD^{WT}$, $RBD^{B.1.351}$, $RBD^{P.1}$ and $RBD^{B.1.1.7}$ present values of -1100 ± 85, -1393 ± 84, -1489 ± 78 and -1013 ± 101 kJ/mol, respectively (**Table S6**). $RBD^{B1.1.7}$ presented binding free energy similar to $RBD^{WT}$, which we have believed that they present insignificant differences in binding affinity. We noted that, in interactions between hACE2 and different $RBD^{WT}$, $RBD^{B.1.351}$, $RBD^{P.1}$ and $RBD^{B.1.1.7}$, Coulomb energy term has a better contribution for free energy than remaining terms, having values of -1380 ± 86, -1453 ± 80, -1461 ± 79 and -1285 ± 76 kJ/mol, respectively (**Table S6**). In addition, Van der Waals energy term presented values of -352 ± 341, -368 ± 33, -430 ± 52 and -297 ± 32 kJ/mol, respectively (**Table S6**). The change in free energy of polar solvation calculated by Poisson-Boltzmann equation (Polar) presented a negative contribution for binding free energy, which presented values of 678 ± 87, 472 ± 85, 454 ± 85 and 612 ± 147 kJ/mol, respectively (**Table S6**). The change in the nonpolar interaction energy under the changes in the protein−solvent accessible surface area (SASA) also were determined e presented values of -46 ± 4, -44 ± 4, -52 ± 6 and -42 ± 6 kJ/mol, respectively (**Table S6**). In general, binding free energy ratio between $RBD^{WT}$ and different $RBD^{B.1.351}$, $RBD^{P.1}$ and $RBD^{B.1.1.7}$ presented values in the range from ~0.9-fold to ~1.4-fold, leading us to believe that the observed reduction between hACE2 and RBD of these SARS-CoV-2 variants presented a moderate effect in decreasing binding affinity. Also, we investigated the contribution of each residue to the binding free energy between RBD and hACE2 (**Figure S7**). Interestingly, most contributions are localized on the hACE2/RBD complex interface and are more frequent in the variants.

**Hydrogen bonding occupancy analysis**

We calculated hydrogen bonding occupancies of residue pairs of each RBD variant/hACE2 interface and compared them with WT (**Table S3**). We noted that there were gains and losses of hydrogen bonding interactions. These results are explained due to different movements of the RBD of these variants, when compared with WT. We observed that residue pairs Q493-$sc^{RBD}$/E35-$sc^{hACE2}$, T449-$sc^{RBD}$/R273-$mc^{hACE2}$, Y449-$sc^{RBD}$/D38-$sc^{hACE2}$ , and Y495-$sc^{RBD}$/E406-$sc^{hACE2}$ kept the hydrogen bonding interactions in hACE2/RBD complex interface, demonstrating that such interactions are relevant in the molecular recognition between hACE2 and RBD. Differently from $RBD^{WT}$, the three variants gained intramolecular hydrogen bonding interaction between (N or Y)501-$sc^{RBD}$/G496-$mc^{RBD}$ but lost between pairs K475-$sc^{RBD}$/E495-$sc^{RBD}$, Y489-$sc^{RBD}$/N487-$sc^{RBD}$ and (K/T/N)417-$sc^{RBD}$/L455-$mc^{RBD}$ (**Table S3**). Interestingly, we also observed that the $RBD^{P.1}$ presents specific hydrogen bonding between pairs Y501-$sc^{RBD}$/Q498-$sc^{RBD}$. Conversely, $RBD^{B.1.1.7}$ presents specific hydrogen bonding interactions between pairs N487-$sc^{RBD}$/A475-$mc^{hACE2}$, T500-$sc^{RBD}$/G354-$mc^{hACE2}$, Y501-$sc^{RBD}$/K353-$mc^{hACE2}$ and Y501-$sc^{RBD}$/Y41-$sc^{hACE2}$. Comparing hydrogen bonding interactions on the interface of the complexes $RBD^{WT}$/hACE2 and $RBD^{P.1}$/hACE2, we also observed this interaction between pairs Q498-$sc^{RBD}$/D38-$sc^{hACE2}$, T500-$sc^{RBD}$/D355-$sc^{hACE2}$ and Y489-$sc^{RBD}$/Y83-$sc^{hACE2}$. Comparing also the complexes $RBD^{WT}$/hACE2 and $RBD^{B.1.1.7}$/hACE2, we also observed hydrogen bonding interactions between pairs

K417-sc$^{RBD}$/D30-sc$^{hACE2}$ and K417-sc$^{RBD}$/H34-sc$^{hACE2}$. When compared with the hACE2/RBD$^{WT}$ complex, we have seen the hydrogen bonding interactions between pairs K31-sc$^{hACE2}$/Q493-sc$^{RBD}$, T500-sc$^{RBD}$/Y41-sc$^{hACE2}$ and T453-sc$^{RBD}$/T449-mc$^{RBD}$ in the RBD$^{B.1.351}$/hACE2 and RBD$^{P.1}$/hACE2 complexes (**Table S3**). Also observed in the B.1.1.7 variant, hydrogen bonding interactions have been seen between R482-sc$^{RBD}$/E489-sc$^{RBD}$ in P.1 and between T453-sc$^{RBD}$ T449-mc$^{RBD}$ in B.1.351 (**Table S3**).

**Cross-correlation analysis**

We calculated the covariance matrix of the spatial displacements of $C_\alpha$ atoms to study internal motions of the hACE2/RBD$^{WT}$ complex and compared them with different SARS-CoV-2 RBD mutants (**Figure S5** and **Tables S4** and **S5**). This computational strategy assumes the hypothesis that residues, even afar, can influence the interactions of other residues. The residue pairs that present absolute correlated and anti-correlated values of more than 0.8 were considered to have more influence over other complex residues (**Table S5**). As shown in **Tables S4-S5** and **Figure S5**, sets of intermolecular and intramolecular covariant pairs changed between different SARS-CoV-2 RBD mutants and RBD$^{WT}$. Both correlated and anticorrelated pairs are affected in E1 and E2 regions of RBD variants, mostly in B.1.1.7 (**Figure S5**). The number of intermolecular covariant pairs for RBD$^{WT}$, RBD$^{B.1.351}$, RBD$^{P.1}$ and RBD$^{B.1.1.7}$ are 21, 8, 135 and 384, respectively (**Table S5**). Among them, only RBD$^{B.1.1.7}$ presented 76 intramolecular negative covariant pairs (**Table S5**). Furthermore, the number of intramolecular positive covariant pairs in hACE2, caused by each RBD, were 3088, 3656, 4160 and 6852, respectively, showing that they increase the internal motions in the host receptor. Conversely, as shown in **Figure S5 e-h** and **Table S5**, the number of intramolecular positive correlations for RBD$^{WT}$, RBD$^{B.1.351}$, RBD$^{P.1}$ and RBD$^{B.1.1.7}$ were 436, 466, 458 and 924, respectively. Interestingly, we did not observe intermolecular anti-correlated pairs between different hACE2/RBD complexes (**Table S5**).

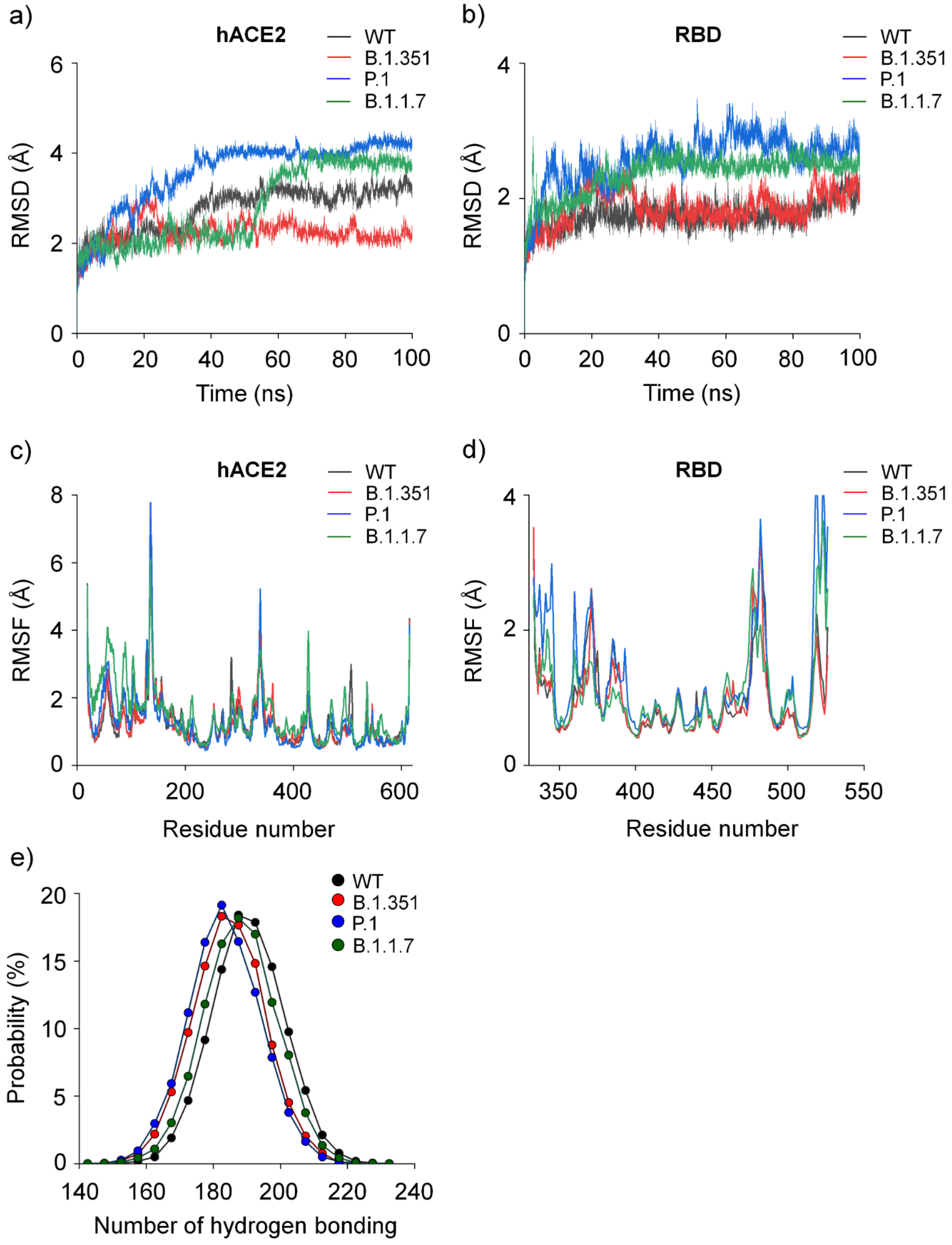


**Figure S1.**

Structural aspects of the SARS-CoV-2 RBD variants and hACE2. **a)** Backbone root-mean-square deviation (RMSD) of hACE2 in complex with RBD (black color) and RBD variants (red, B.1.351; blue, P.1; green, B.1.1.7). **b)** Backbone RMSD of

RBD (black color) and RBD variants (red, B.1.351; blue, P.1; green, B.1.1.7). **c)** Backbone root-mean-square fluctuation (RMSF) of hACE2 in complex with RBD (black color) and RBD variants (red, B.1.351; blue, P.1; green, B.1.1.7). **d)** Backbone RMSF of RBD (black color) and RBD variants (red, B.1.351; blue, P.1; green, B.1.1.7). **e)** Probability distributions of the number of hydrogen bonding interactions of RBD (black color) and its variants (red, B.1.351; blue, P.1; green, B.1.1.7) in complex with hACE2.

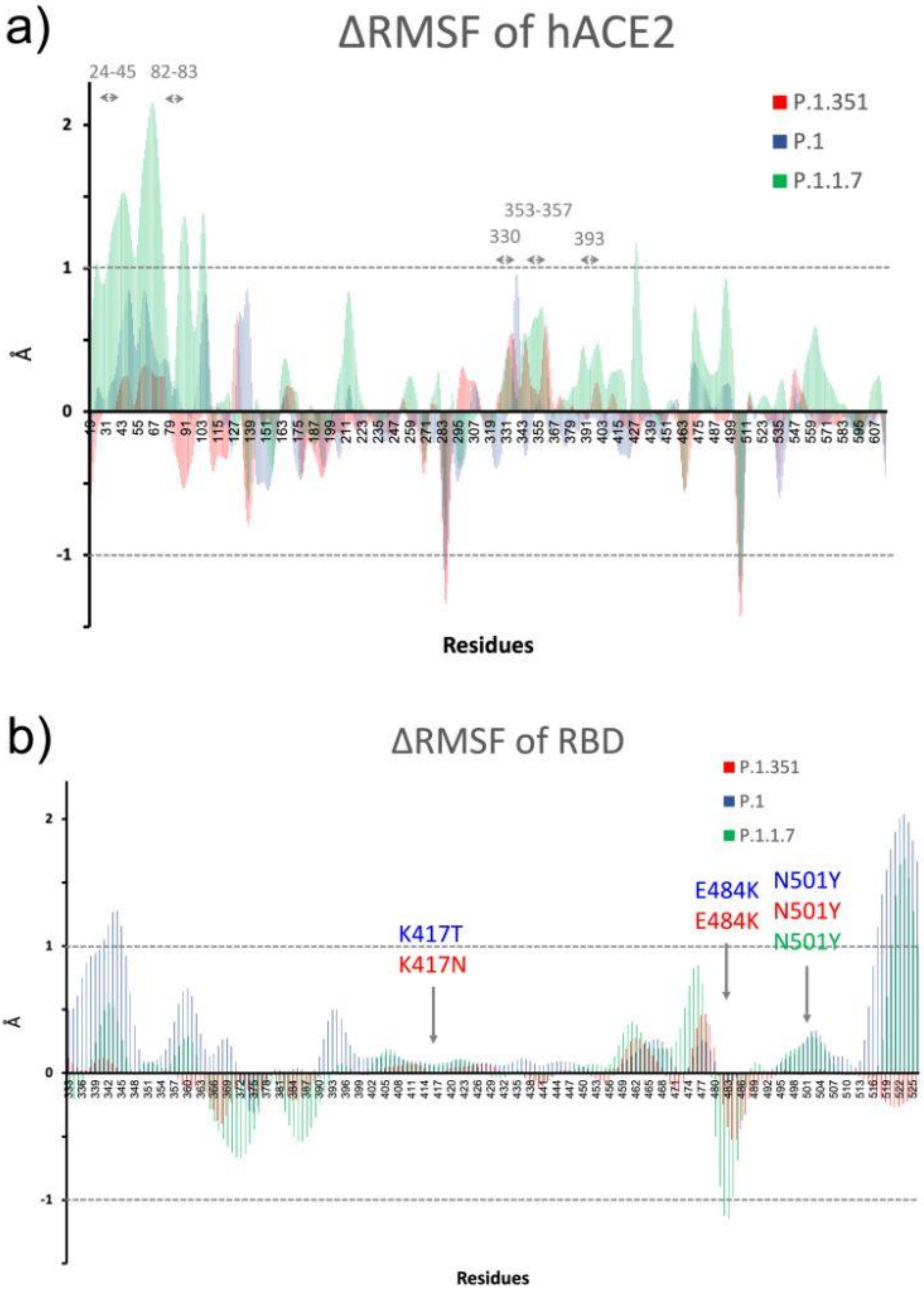


**Figure S2. Backbone root-mean-square fluctuation differences of the hACE2 interacting with RBD variants. a)** Relative backbone RMSF$^{hACE2}$. **b)** Relative backbone RMSF$^{RBD}$. Both panels **a-b** adopt hACE2/RBD$^{WT}$ complex as reference.

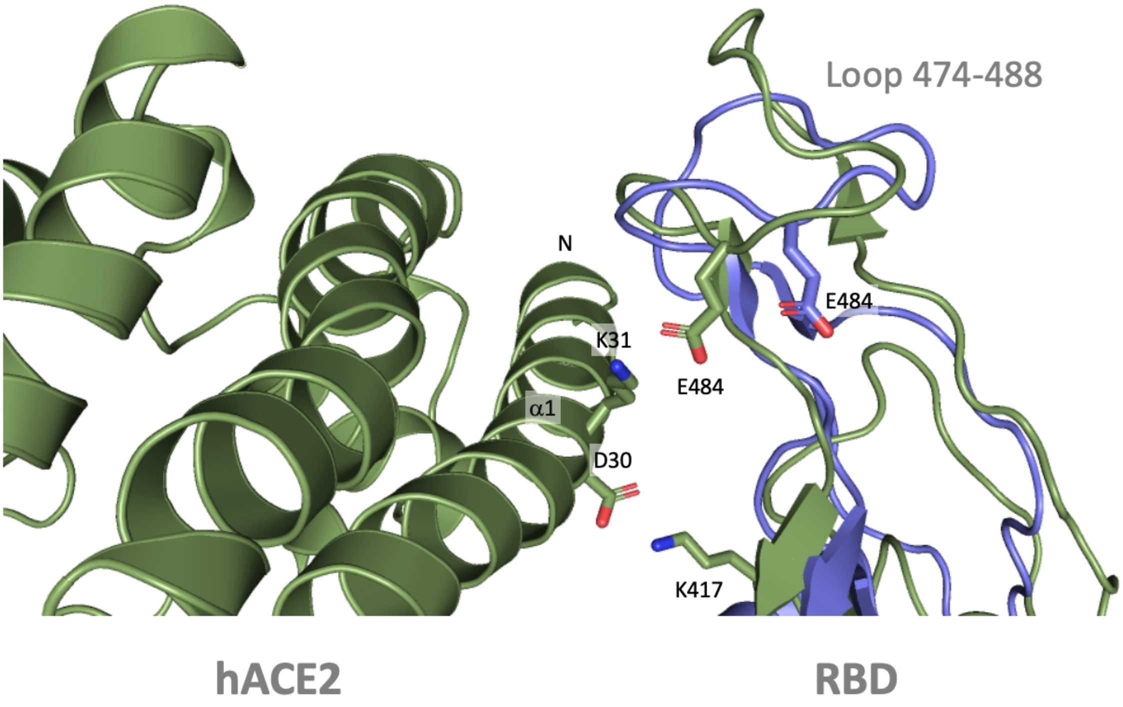


**Figure S3.** Comparison between modeled RBD$^{B.1.1.7}$ (green color) with trimeric Spike RBD$^{B.1.1.7}$ obtained by cryo-electron microscopy (blue color, PDB ID 7MJG [17]).

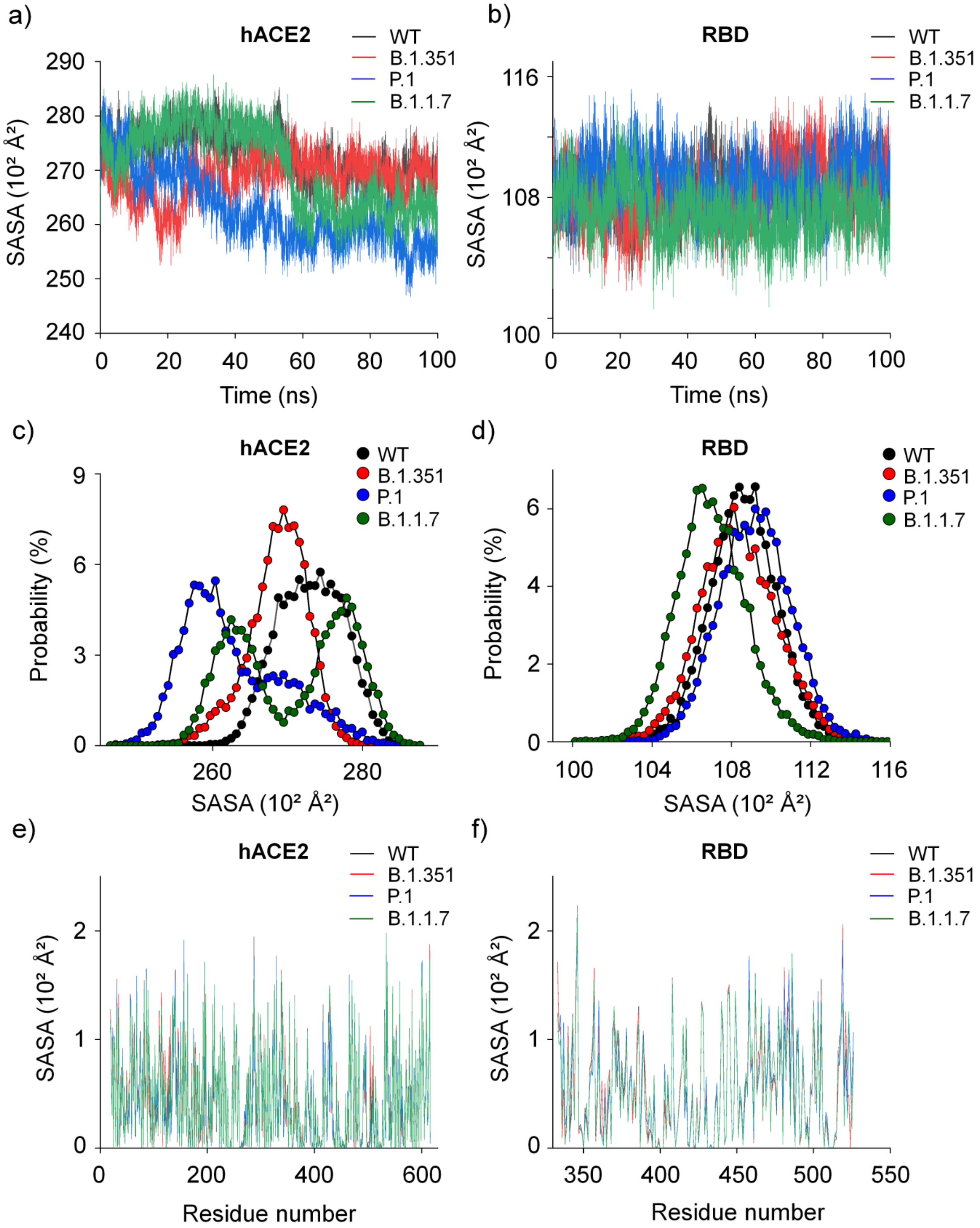


**Figure S4.** Solvent-accessible surface area (SASA) information of SARS-CoV-2 RBD variants and hACE2. SASA of **a)** hACE2 while interacting with **b)** RBD$^{WT}$ and its variants. **c)** hACE2 and **d)** RBD SASA probability distributions. SASA per residue of **e)** hACE2 in complex with **f)** RBD (black color) and RBD variants (red, B.1.351; blue, P.1; green, B.1.1.7). RBD$^{WT}$ and its variants did not present significant differences in SASA along MD trajectory. However, in hACE2, SASA values presented changes and such alterations are associated with conformational states of hACE2 due to closing of its active site mediated by RBD and its variants.

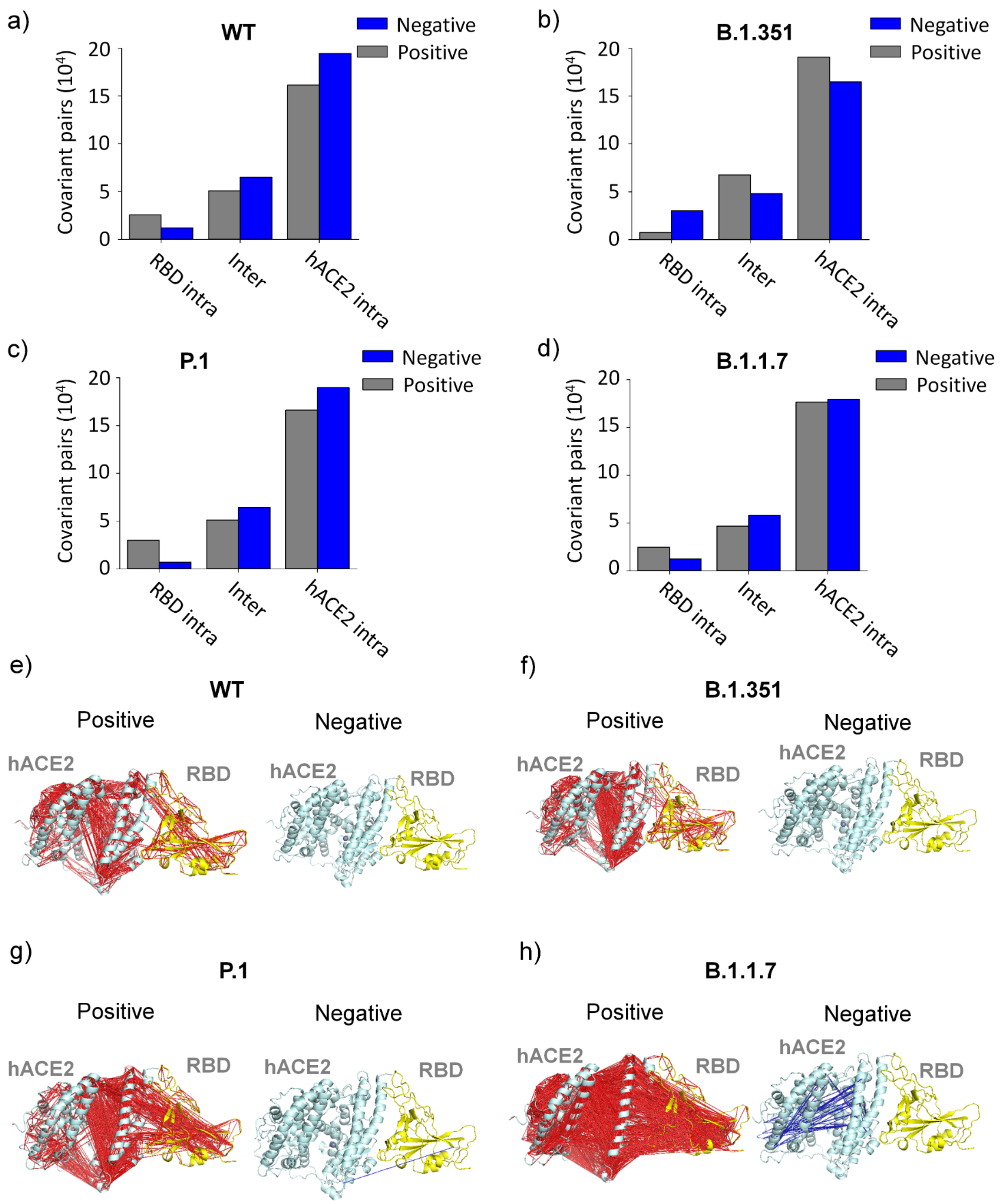


**Figure S5.** Cross-correlation analysis between SARS-CoV-2 RBD variants and hACE2. Covariance analysis of MD trajectories of RBD variants and WT. panels **a-d** Number of covariant pairs observed in the interaction of $RBD^{WT}$ and their variants with hACE2. Panels **e-h** Pairs of $C_{\alpha}$ exhibiting covariance higher than 0.8 were analyzed in the interaction of RBD$^{WT}$ and their variants with hACE2. The red and blue colors represent positive and negative correlations, respectively.

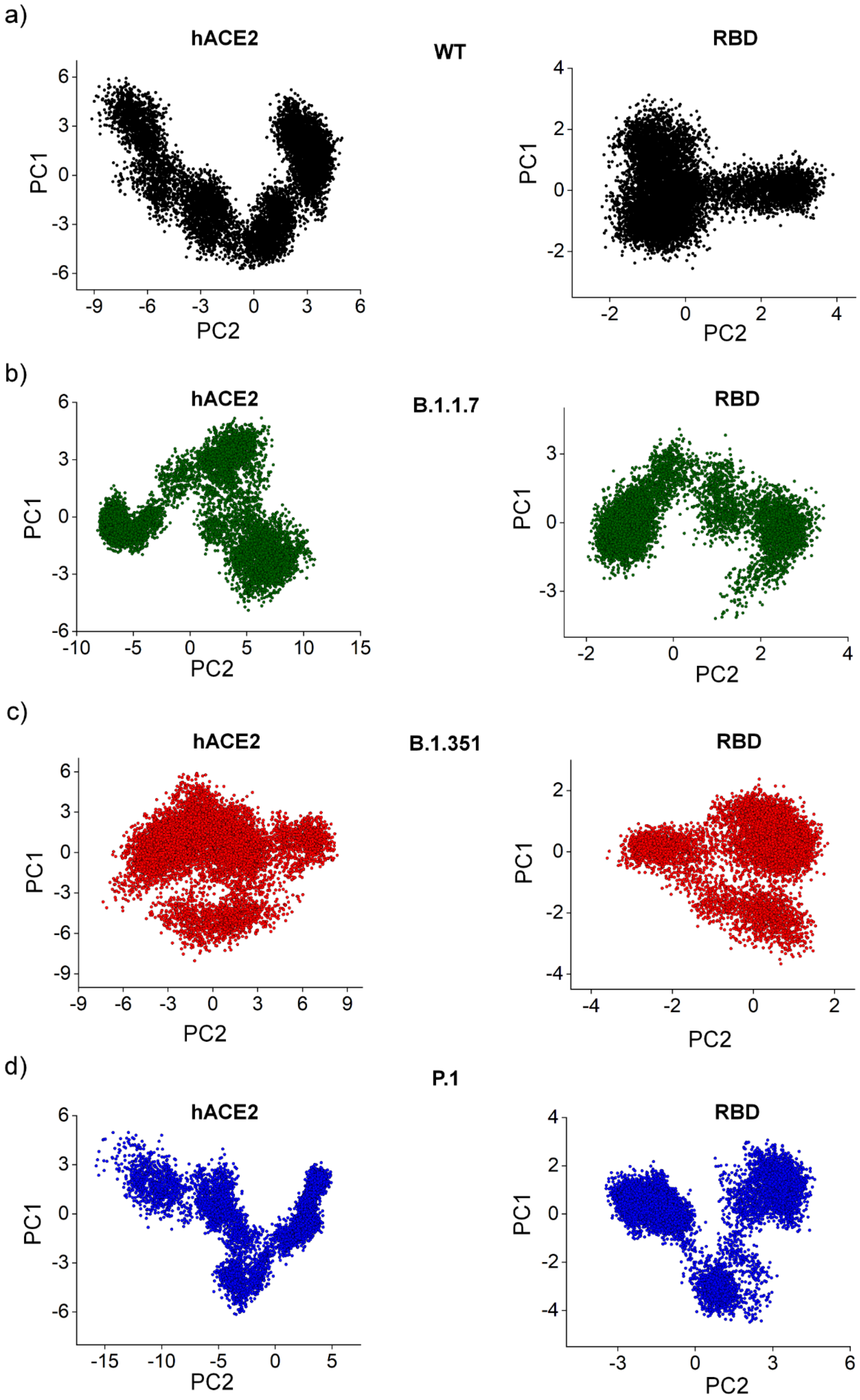


**Figure S6.** Molecular dynamics trajectories obtained by principal component analysis. Using first and second components, painels a-d show trajectory scores of hACE2 while interacting with **a)** RBD$^{WT}$, **b)** RBD$^{B.1.1.7}$, **c)** RBD$^{B.1.351}$, and **d)** RBD$^{P.1}$.

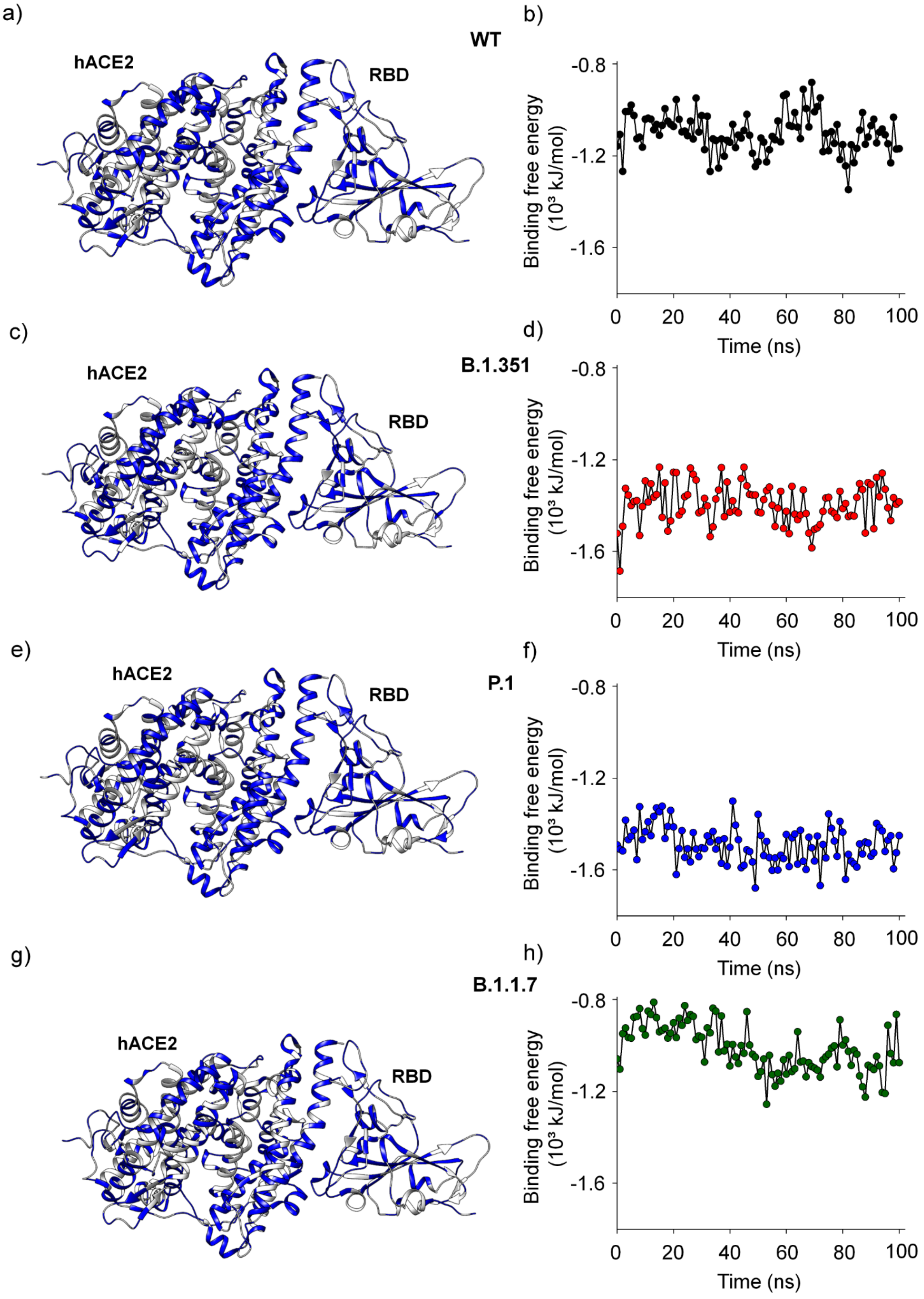


**Figure S7.** Binding free energies of RBD and its variants for formation of a complex with hACE2. Painels **b)**, **d)**, **f)** and **h)** show binding free energies as function of time using MD trajectories of the $RBD^{WT}$ and its variants in complex with hACE2. Painels

**a)**, **c)**, **e)** and **g)** show energy contributions of each residue for binding free energy, in which blue and white colors represent favorable and unfavorable contributions, respectively.

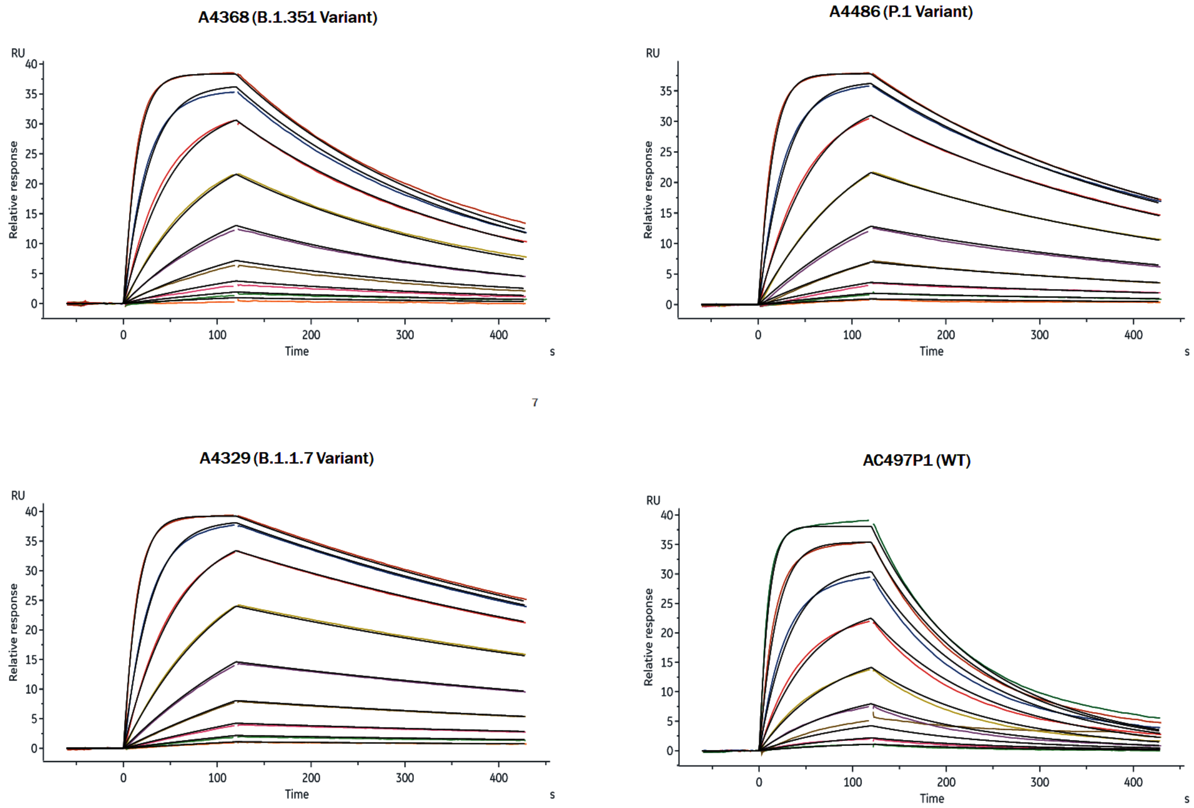


**Figure S8.** Surface plasmon resonance (SPR) data using immobilized recombinant dimer hACE2 and different concentrations of purified recombinant Spike RBD and their variants, as described in materials and methods. The experimental data were fitted to a 1:1 binding model using Biacore Evaluation Software (GE Healthcare) to calculate the kinetics parameters ($k_a$ and $k_d$) shown in **Table 1** and **Table S7**. At least eight different RBD concentrations were used to calculate the $K_D$ values.

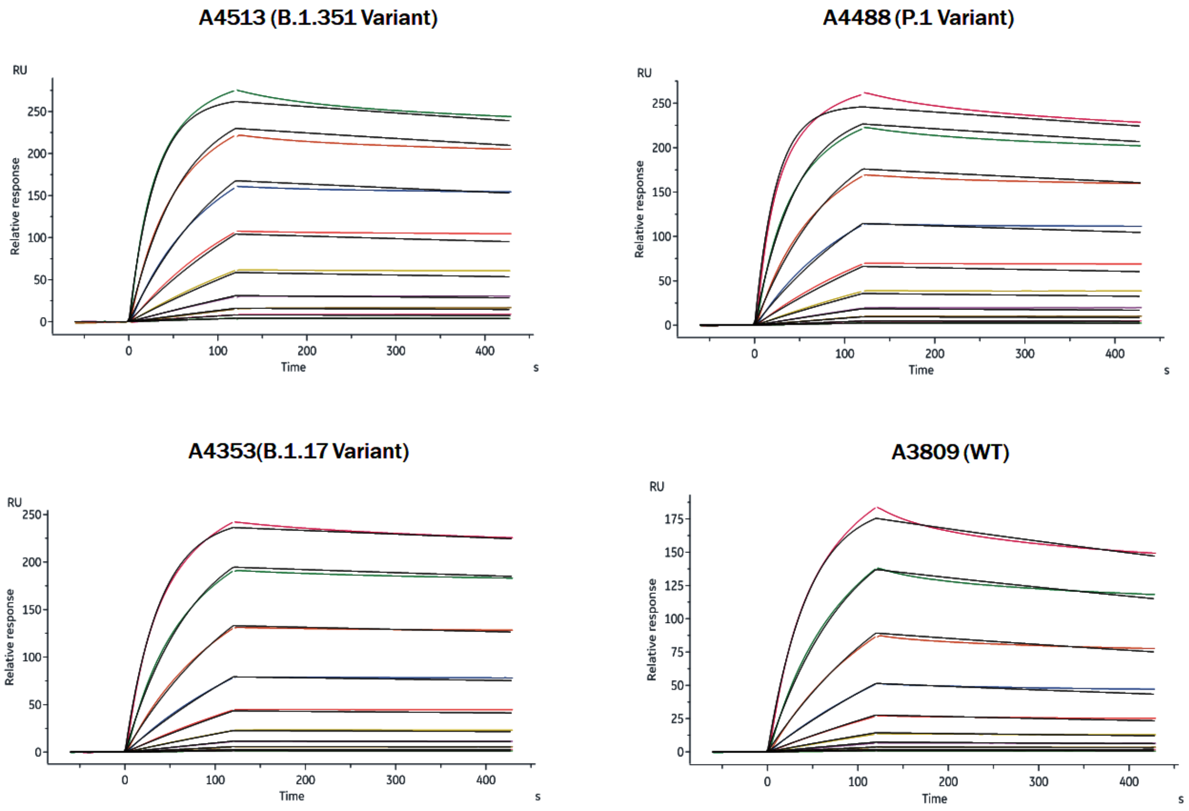


**Figure S9.** Surface plasmon resonance (SPR) data using immobilized recombinant dimer hACE2 and different concentrations of purified recombinant Spike trimer and their variants, as described in materials and methods. The experimental data were fitted to a 1:1 binding model using Biacore Evaluation Software (GE Healthcare) to calculate the kinetics parameters ($k_a$ and $k_d$) shown in **Table 1** and **Table S7**. At least seven different Spike concentrations were used to calculate the $K_D$ values.

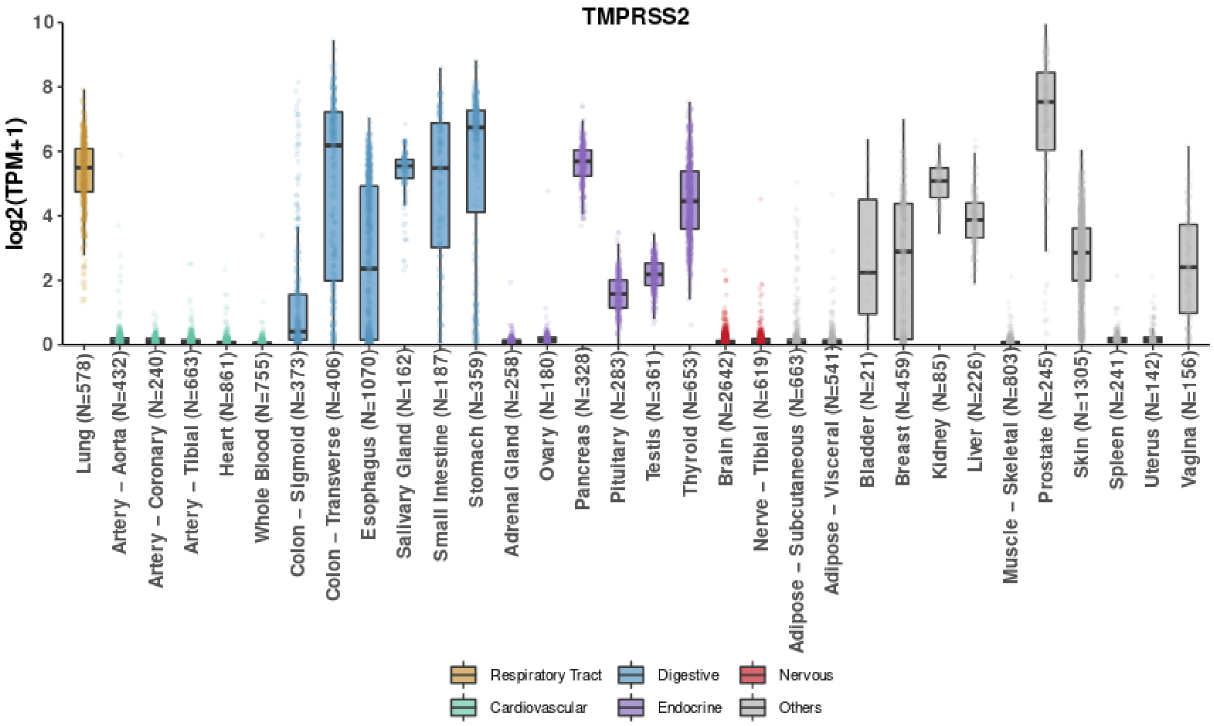


**Figure S10.** TMPRSS2 is mainly expressed in digestive, endocrine and few other human tissues or organs. Box colors represent different tissue groups according to their major functions. Numbers of samples are shown in parentheses.

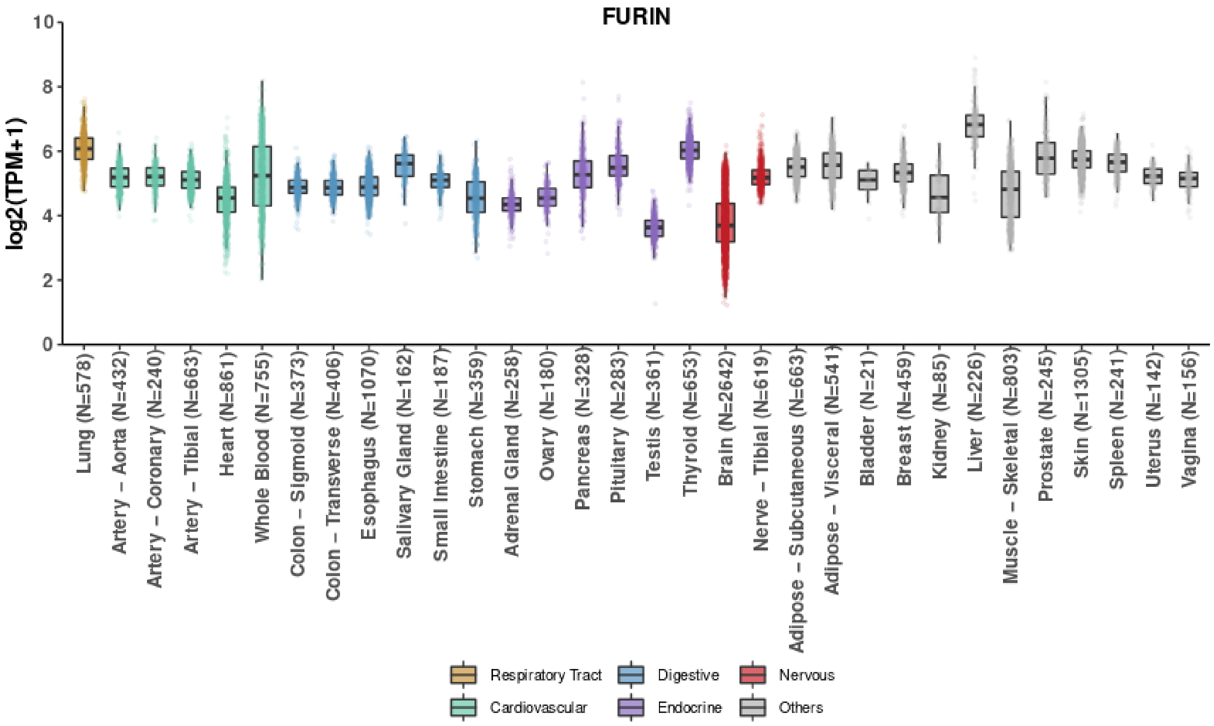


**Figure S11.** FURIN is highly expressed in all human tissues. Box colors represent different tissue groups according to their major functions. Numbers of samples are shown in parentheses.

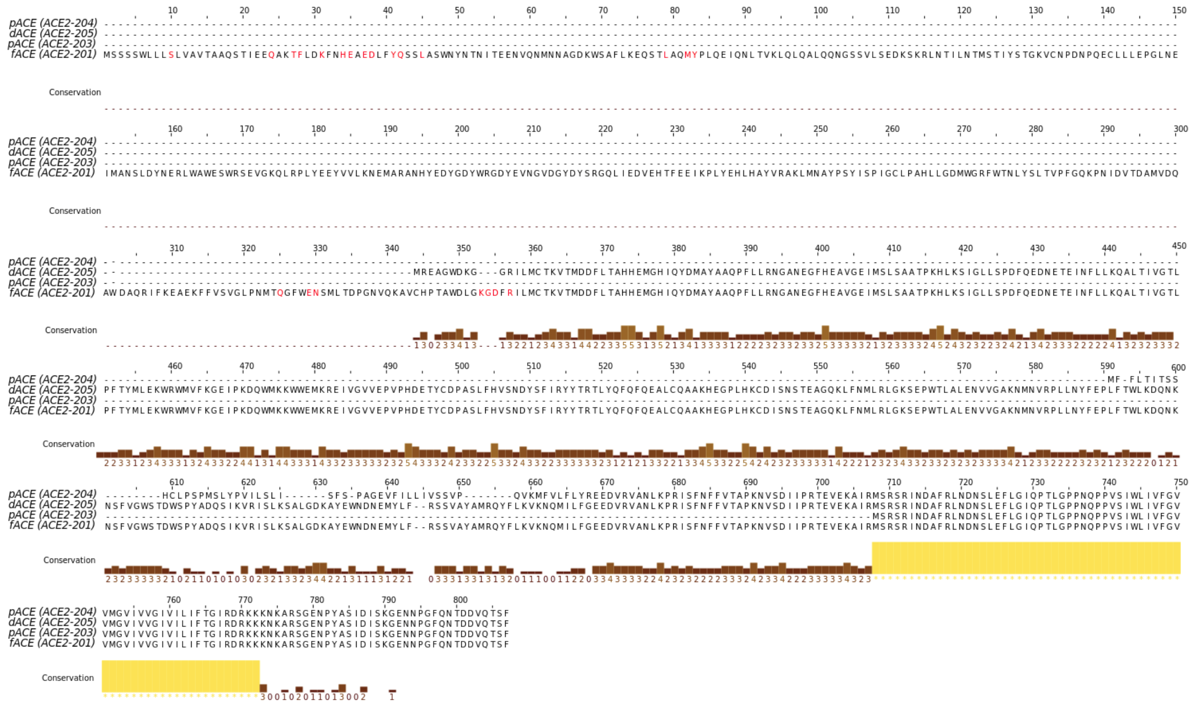


**Figure S12.** Multiple alignment of translated ACE2 transcripts. Transcripts were classified by fACE (full-length ACE), which is represented by the canonical isoform ACE2-201 and have all functional domains; dACE, that lacks RBD domain interaction residues; and pACE, representing all other shorter ACE2 isoforms not neither reported in fACE or dACE classes. Residues marked in red correspond to direct contact portions of hACE2 with RBD domain.

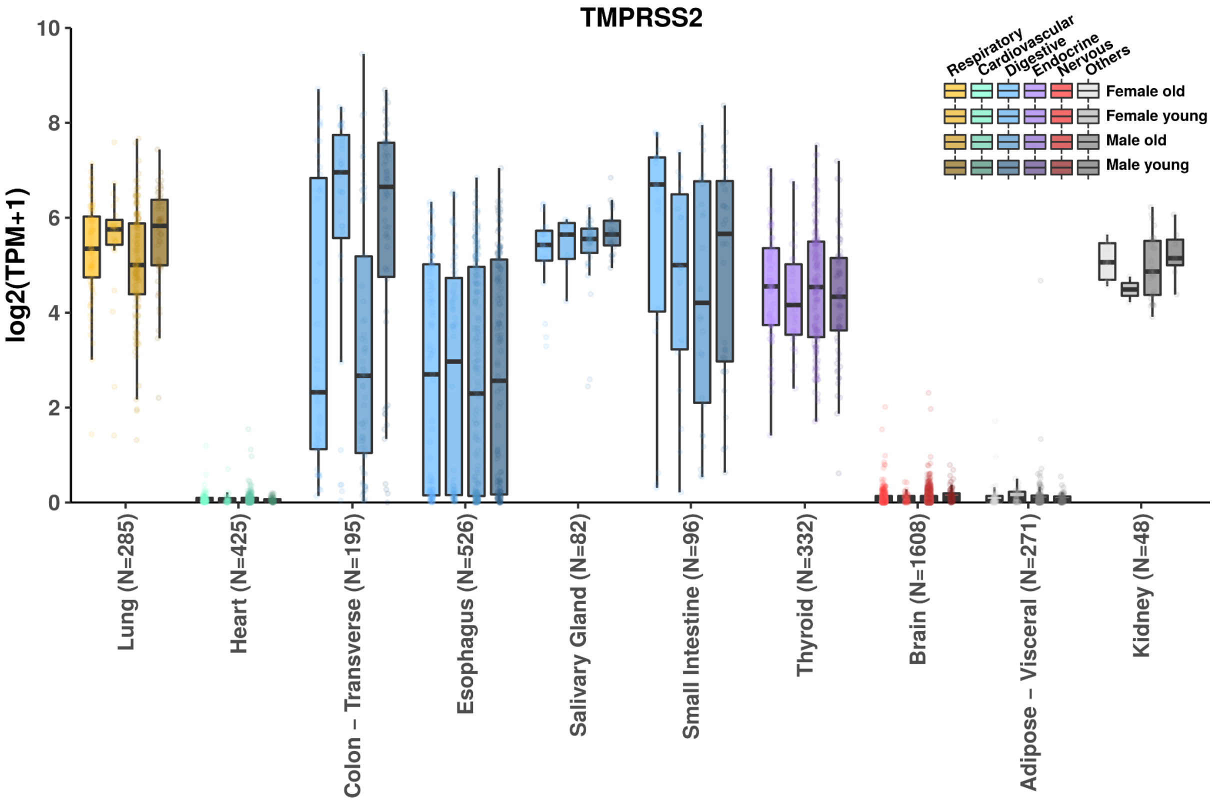


**Figure S13.** Tissue-wide TMPRSS2 expression by age and gender. Box colors represent different tissue groups according to their major functions. Box shades refer to distinct clinical variables, either age (lighter) or gender (darker).

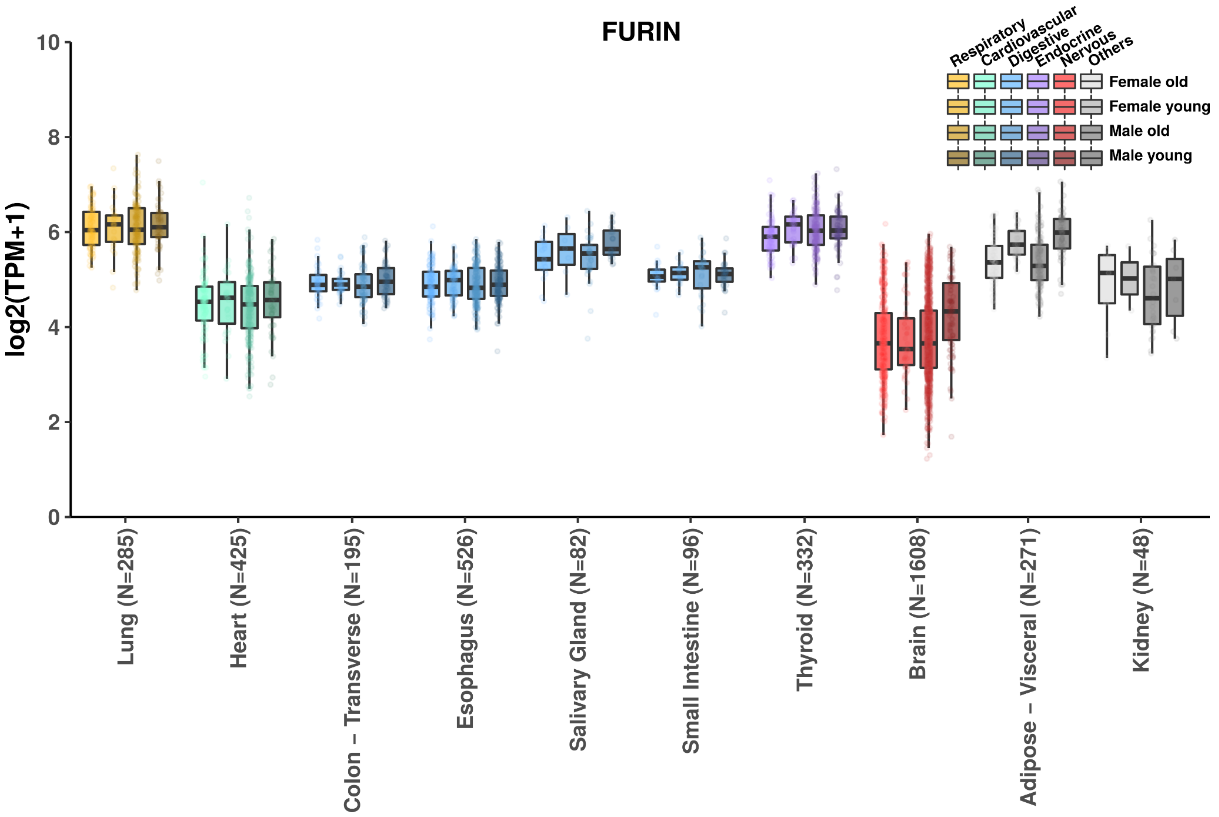


**Figure S14.** Tissue-wide FURIN expression by age and gender. Box colors represent different tissue groups according to their major functions. Box shades refer to distinct clinical variables, either age (lighter) or gender (darker).

**Table S1.**

Insights about SARS-CoV-2 variants. Insights about origin, mutations and number of countries relating cases associated with each variant. NA, not available.

| SARS-CoV-2 Variant | Country First Detected | Number of countries that reported the variant (25 May 2021) | Spike Protein mutations |
|---|---|---|---|
| B.1.1.7[18] (Alpha) | United Kingdom, London | 149[19] | Deletion 69/70<br>Deletion 144/145<br>N501Y<br>A570D<br>D614G<br>P681H<br>T716I<br>S982A<br>D1118H |
| B.1.351[20] (Beta) | South Africa | 102[19] | L18F<br>D80A<br>D215G<br>Deletion 242/244<br>R246I<br>K417N<br>E484K<br>N501Y<br>D614G<br>A701V |
| P.1[21] (Gamma) | Brazil, Manaus | 59[19] | L18F<br>T20N<br>P26S<br>D138Y<br>R190S<br>K417T<br>E484K<br>N501Y<br>D614G<br>H655Y<br>T1027I<br>V1176F |
| B.1.617.2[22] (Delta) | India | 54[19] | T19R<br>Deletion 157/158<br>L452R<br>T478K<br>D614G<br>P681R<br>D950N |
| B.1.427/429[23,24] (Epsilon) | United States, California | 29 (01 Abril 2021)[25] | S13I<br>W152C |

| | | | |
|---|---|---|---|
| | | | L452R<br>D614G |
| B.1.526[26]<br>(Iota) | United States,<br>New York | NA | L5F<br>T95I<br>D253G<br>E484K<br>D614G<br>A701V |
| B.1.617.1[27]<br>(kappa) | India | 41[19] | L452R<br>E484Q<br>D614G<br>P681R<br>Q1071H |

**Table S2.** Insights about SARS-CoV-2 variants. Compared with wild-type, variants presented significant increases of transmissibility, symptoms, time of infectivity, viral load, severity, lethality, binding affinity and replication rate. ND: undetermined information; NA: unavailable information.

| | **B.1.1.7** | **P.1** | **B.1.351** | **B.1.429/427** |
|---|---|---|---|---|
| **Increase of transmissibility** | 10 - 75% [28,29,30,31,32,33,34,35,36] | 12 - 46% [28,37,35] | 55%[35,36] | 6-24%[38,31] |
| **Higher viral load** | 20 - 30% [39,40,41,16,17,42,35] | 10 fold [43,35] | ND [40,35] | 2 fold [38] |
| **More severe disease** | 40-65%[36,44,45] | ND[45] | ND[45] | NA |
| **Higher lethality** | 8 - 71% [46,30,47,48,49,36,44] | ND [45] | ND [45,50] | NA |
| **More binding affinity to ACE2 in relation to $RBD^{WT}$** | ~2 fold[51] | NA | ~4.6 fold[51] | NA |

**Table S3.** Hydrogen bonding occupancy (%) of interactions between RBD of SARS-CoV-2 variants and hACE2. Hydrogen bond occupancy (%) analysis of the RBD residues and its variants that make contact with hACE2 along different MD trajectories. Table shows intramolecular and intermolecular interactions in hACE2 and RBD. Cut-off: hydrogen bond distance of 3 Å between hydrogen and nitrogen or oxygen. sc = side chain and mc = main chain.

| **HB donor** | **HB acceptor** | **WT** | **B.1.351** | **P.1** | **B.1.1.7** |
|---|---|---|---|---|---|

| | | | | | |
|---|---|---|---|---|---|
| R482-sc$^{RBD}$ | E489-sc$^{RBD}$ | | | 90.7 | 87.7 |
| N487-sc$^{RBD}$ | A475-mc$^{hACE2}$ | | | | 15.8 |
| Q493-sc$^{RBD}$ | E35-sc$^{hACE2}$ | 51.9 | 40.4 | 42.8 | 4.4 |
| Q498-sc$^{RBD}$ | D38-sc$^{hACE2}$ | 38.7 | | 14.6 | |
| E489-mc$^{RBD}$ | E489-sc$^{RBD}$ | 50.0 | 54.9 | 33.5 | 50.1 |
| K31-sc$^{hACE2}$ | Q493-sc$^{RBD}$ | 19.6 | 9.0 | 16.7 | |
| K475-sc$^{RBD}$ | E495-sc$^{RBD}$ | 23.4 | | | |
| T449-sc$^{RBD}$ | R273-mc$^{hACE2}$ | 26.9 | 41.3 | 57.3 | 27.9 |
| T453-sc$^{RBD}$ | T449-mc$^{RBD}$ | 44.2 | | 41.0 | |
| T500-scRBD | Y41-sc$^{hACE2}$ | 35.8 | 10.3 | 13.7 | |
| T500-sc$^{RBD}$ | D355-sc$^{hACE2}$ | 1.7 | | 19.0 | |
| T500-sc$^{RBD}$ | G354-mc$^{hACE2}$ | | | | 33.1 |
| Y449-sc$^{RBD}$ | D38-sc$^{hACE2}$ | 63.2 | 59.2 | 56.7 | 4.7 |
| T453-sc$^{RBD}$ | T449-mc$^{RBD}$ | | 33.7 | | 45.8 |
| K484-sc$^{RBD}$ | E75-sc$^{hACE2}$ | | 1.5 | 1.2 | |
| F490-mc$^{RBD}$ | E484-sc$^{RBD}$ | 0.1 | | | 66.5 |
| G485-mc$^{RBD}$ | E484-sc$^{RBD}$ | 17.0 | | | |
| K31-sc$^{hACE2}$ | E484-sc$^{RBD}$ | ~0.0 | | | 47.4 |
| F486-mc$^{RBD}$ | E484-sc$^{RBD}$ | 19.0 | | | |
| Y489-sc$^{RBD}$ | N487-sc$^{RBD}$ | 15.4 | | | |
| Y489-sc$^{RBD}$ | Y83-sc$^{hACE2}$ | 4.1 | | 22.5 | |
| Y495-sc$^{RBD}$ | E406-sc$^{hACE2}$ | 59.2 | 26.5 | 15.5 | 28.9 |
| K417-sc$^{RBD}$ | D30-sc$^{hACE2}$ | 51.5 | | | 28.2 |
| K417-sc$^{RBD}$ | H34-sc$^{hACE2}$ | 13.5 | | | 8.2 |
| K417-sc$^{RBD}$ | L455-mc$^{RBD}$ | 5.4 | | | |
| Y501-sc$^{RBD}$ | G496-mc$^{RBD}$ | | 21.3 | 13.9 | 1.7 |
| Y501-sc$^{RBD}$ | Q498-sc$^{RBD}$ | | | 25.9 | |
| Y501-sc$^{RBD}$ | K353-mc$^{hACE2}$ | | | | 18.7 |
| Y501-sc$^{RBD}$ | Y41-sc$^{hACE2}$ | | | | 2.8 |

**Table S4.** Count of positive and negative covariants using cross-correlation analysis. Covariance analysis of MD trajectories of RBD variants and WT. Total covariant pairs observed in the interaction of RBD$^{WT}$ and its variants with hACE2.

| | Correlation | Intra-RBD | Inter | Intra-ACE2 |
|---|---|---|---|---|
| **WT** | positive | 25488 | 50848 | 161492 |
| | negative | 11954 | 64970 | 194320 |
| **B.1.351** | positive | 7300 | 67747 | 190816 |
| | negative | 30142 | 48071 | 164996 |
| **P.1** | positive | 30200 | 51365 | 166086 |
| | negative | 7242 | 64453 | 189726 |
| **B.1.1.7** | positive | 24822 | 46852 | 176502 |

| | | | | |
|---|---|---|---|---|
| | negative | 12620 | 58220 | 179310 |

**Table S5.** Count of relevant positive and negative covariants using cross-correlation analysis. Pairs of $C_{\alpha}$ exhibiting covariance higher than 0.8 were analyzed in the interaction of $RBD^{WT}$ and its variants while interacting with hACE2.

| | Correlation | Intra-RBD | Inter | Intra-ACE2 |
|---|---|---|---|---|
| **WT** | positive | 436 | 21 | 3088 |
| | negative | 0 | 0 | 0 |
| **B.1.351** | positive | 466 | 8 | 3656 |
| | negative | 0 | 0 | 0 |
| **P.1** | positive | 458 | 135 | 4160 |
| | negative | 0 | 1 | 0 |
| **B.1.1.7** | positive | 924 | 384 | 6852 |
| | negative | 0 | 0 | 76 |

**Table S6.** Binding free energy between RBD of different SARS-CoV-2 variants and hACE2.

| | Energy (kJ/mol) | | | | |
|---|---|---|---|---|---|
| **Variants** | **vdW** | **Coul** | **Polar** | **SASA** | **ΔG** |
| B.1.351 | -368 +/- 33 | -1453 +/- 80 | 472 +/- 85 | -44 +/- 4 | -1393 +/- 84 |
| P.1 | -430 +/- 52 | -1461 +/- 79 | 454 +/- 85 | -52 +/- 6 | -1489 +/- 78 |
| B.1.1.7 | -297 +/- 32 | -1285 +/- 76 | 612 +/- 147 | -42 +/- 6 | -1013 +/- 101 |
| WT | -352 +/- 341 | -1380 +/- 86 | 678 +/- 87 | -46 +/- 4 | -1100 +/- 85 |

**Table S7.** Kinetic parameters obtained from surface plasmon resonance (SPR) assays.

| Variant | Capture Level (RU) | Spike Protein Concentrations (nM) | $k_a$ ($10^6$ $M^{-1}s^{-1}$) | $k_d$ ($10^{-4}$ $s^{-1}$) | $K_D$ (nM) | $R_{max}$ (RU) | $Chi^2$ ($RU^2$) |
|---|---|---|---|---|---|---|---|
| $RBD^{B.1.351}$ | 173.3 | 0.244-62.5 | 1.2 | 39.4 | 3.3 | 40.3 | 0.17 |
| $RBD^{P.1}$ | 170.5 | 0.244-62.5 | 1.3 | 28.8 | 2.2 | 39.1 | 0.06 |
| $RBD^{B.1.1.7}$ | 174.1 | 0.244-62.5 | 1.3 | 15.5 | 1.2 | 40.0 | 0.04 |
| $RBD^{WT}$ | 176.1 | 0.488-125 | 0.9 | 91.6 | 10.3 | 41.2 | 0.61 |
| Spike $trimer^{B.1.351}$ | 178.4 | 0.488-125 | 0.3 | 3.0 | 1.1 | 269.0 | 15.98 |
| Spike $trimer^{B.1.1.7}$ | 174.5 | 0.488-250 | 0.1 | 1.7 | 1.6 | 248.8 | 2.35 |
| Spike $trimer^{P.1}$ | 170.5 | 0.488-250 | 0.2 | 3.0 | 1.8 | 249.3 | 15.22 |

| | | | | | | |
|---|---|---|---|---|---|---|
| S trimer$^{WT}$ | 169.4 | 0.488-250 | 0.1 | 5.8 | 6.4 | 192.8 | 1.53 |

**Table S8.** Experimental $K_D$ obtained by different kinetic assays. ND: not determined.

| Experimental assay | Construction | $K_D^{B.1.351}$ (nM) | $K_D^{B.1.1.7}$ (nM) | $K_D^{P.1}$ (nM) | $K_D^{WT}$ (nM) |
|---|---|---|---|---|---|
| Microscale thermophoresis[51] | RBD | 87.6 | 203.7 | ND | 402.5 |
| bio-layer interferometry[52] | RBD | ND | 22 | ND | 133 |
| SPR[53] | RBD | 48.2 | 13.1 | ND | 56.9 |
| SPR[54] | RBD | 4.8 | 9.1 | ND | 14 |
| bio-layer interferometry[55] | Spike trimer | 3.2-13.7 | 0.1-2.4 | ND | ND |
| SPR[56–58] | RBD | ND | ND | ND | 5.1 - 44.2 |

**Table S9.** Eigenvalues of the principal component 1 and 2, PC1 and PC2 respectively, obtained from diagonalizations of the 7119x7119 covariance matrix (representing backbone hACE2) and 5373x5373 covariance matrix (representing backbone RBD).

| | Protein | Eigenvalue (PC1) | Eigenvalue (PC2) |
|---|---|---|---|
| WT | hACE2 | 12.9 | 6.4 |
| | RBD | 1.7 | 0.9 |
| B.1.1.7 | hACE2 | 32.5 | 4.0 |
| | RBD | 2.3 | 1.0 |
| B.1.351 | hACE2 | 9.5 | 5.8 |
| | RBD | 1.3 | 1.0 |
| P.1 | hACE2 | 20.3 | 3.3 |
| | RBD | 4.6 | 2.1 |
